# Evolution of stickleback spines through independent *cis*-regulatory changes at *HOXDB*

**DOI:** 10.1101/2021.12.14.472698

**Authors:** Julia I. Wucherpfennig, Timothy R. Howes, Jessica N. Au, Eric H. Au, Garrett A. Roberts Kingman, Shannon D. Brady, Amy L. Herbert, Thomas E. Reimchen, Michael A. Bell, Craig B. Lowe, Anne C. Dalziel, David M. Kingsley

## Abstract

Understanding the genetic mechanisms leading to new traits is a fundamental goal of evolutionary biology. We show that *HOXDB* regulatory changes have been used repeatedly in different stickleback fish species to alter the length and number of bony dorsal spines. In *Gasterosteus aculeatus*, a variant *HOXDB* allele is genetically linked to shortening an existing spine and adding a spine. In *Apeltes quadracus*, a variant allele is associated with lengthening an existing spine and adding a spine. The alleles alter the same conserved non-coding *HOXDB* enhancer by diverse molecular mechanisms, including SNPs, deletions, and transposable element insertions. The independent *cis*-acting regulatory changes are linked to anterior expansion or contraction of *HOXDB* expression. Our findings support the long-standing hypothesis that natural *Hox* gene variation underlies key morphological patterning changes in wild populations and illustrate how different mutational mechanisms affecting the same region may produce opposite gene expression changes with similar phenotypic outcomes.

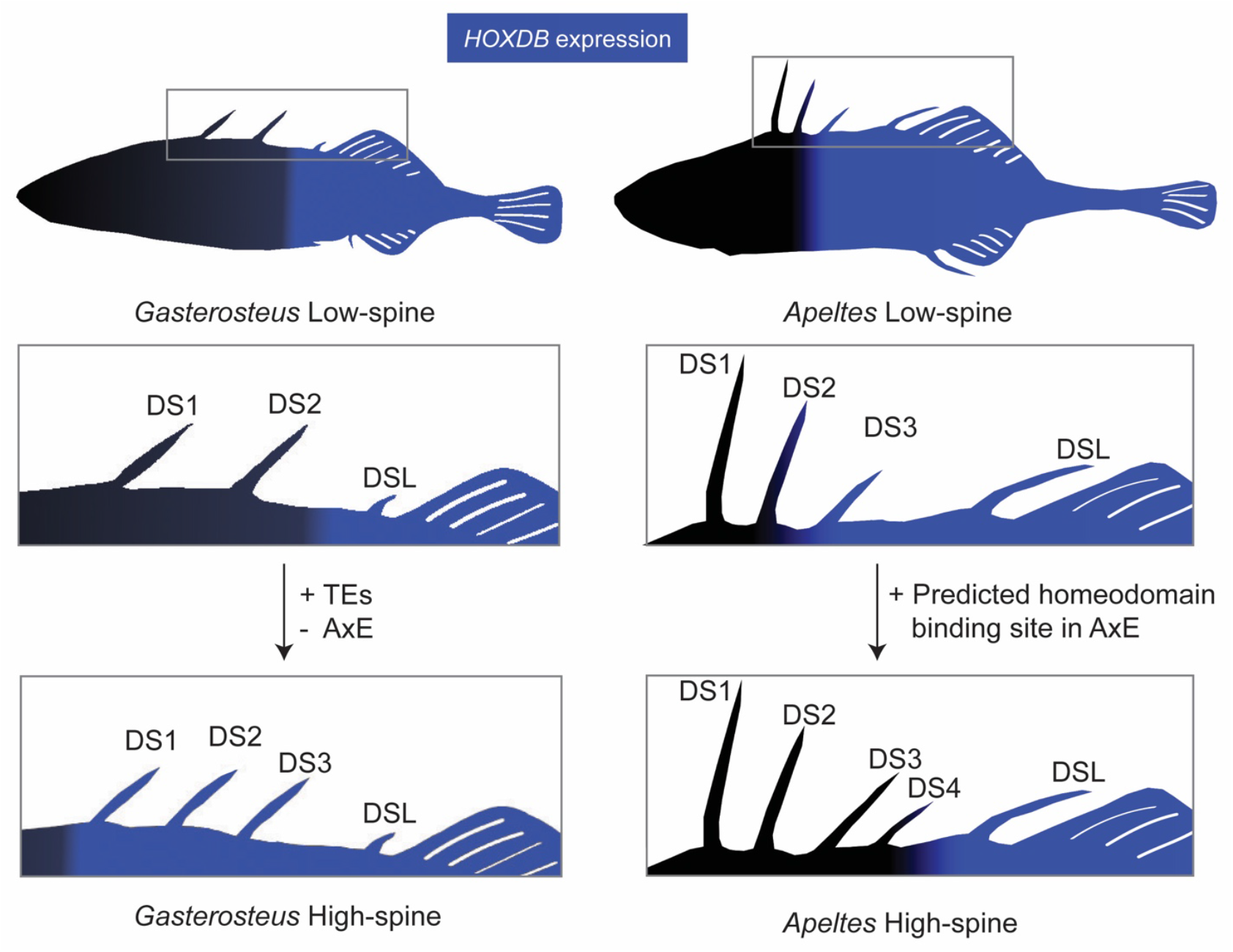

## Introduction

The origins of diverse vertebrate body plans have fascinated comparative anatomists and evolutionary biologists for centuries (Darwin, 1859; Owen, 1848). Although studies over the last forty years have now identified many key cellular pathways required for normal body axis formation and development using induced mutations in model organisms, it remains challenging to identify specific changes in genes and regulatory regions that underlie the diversity of body forms and traits in wild species (Stern and Orgogozo, 2008, 2009).

*Hox* genes were one of the first classes of major developmental genes to be identified and analyzed in comparative studies across many animal groups. They were initially discovered by linked clusters of mutations in *Drosophila* that had the remarkable ability to transform particular body segments into others (Lewis, 1978). Molecular cloning studies revealed that *Hox* loci consist of clustered homeodomain transcription factor genes, whose expression pattern along the Anterior-Posterior (A-P) body axis was correlated with their physical position along the chromosome (Bender et al., 1983; Carroll et al., 2005; Harding et al., 1985; Izpisua-Belmonte et al., 1991; Scott and Weiner, 1984).

In an early review of genetic work on homeotic loci, Ed Lewis hypothesized that regulatory mutations in *Hox* genes might underlie classic A-P patterning differences between species, such as four-winged versus two-winged insects (Lewis, 1978). Although subsequent studies showed that *Hox* expression patterns are actually conserved between two-winged fruit flies and four-winged butterflies (Carroll et al., 1995; Warren et al., 1994), the important role of *Hox* genes in controlling many aspects of body patterning has led to speculation that mutations in these genes underlie key morphological differences in nature (Carroll et al., 2005; Goldschmidt, 1940). Variation in *Hox* cluster number and structure across different taxa support this idea, and intriguing correlations can be drawn between morphological differences in body traits and *Hox* expression changes in other animal groups (Averof and Patel, 1997; Burke et al., 1995; Carroll et al., 2005). On the other hand, much of the diversification and expansion of *Hox* clusters occurred prior to well-known morphological changes among animal phyla (Carroll, 1995). Furthermore, many laboratory mutations in *Hox* genes lead to reduced viability or fertility, and prominent evolutionary biologists (Liu et al., 2019; Mayr, 1970), as well as critics of evolutionary biology (Wells, 2000), have suggested that natural mutations in *Hox* genes would most likely lead to “hopeless monsters” rather than adaptive changes in wild species. Natural differences in leg trichomes and abdominal pigmentation have previously been linked to genetic variation in *Hox* loci in insects, with regulatory mutations providing a possible mechanism for bypassing the broader deleterious consequences seen with many laboratory mutations (Stern, 1998; Tian et al., 2019). However, few detailed examples exist for the long-postulated idea that genetic changes in *Hox* loci may also be the basis for major changes in skeletal structures along the A-P body axis of wild vertebrates (Burke et al., 1995; Shashikant et al., 1998).

Almost a third of extant vertebrate species fall in the large and diverse Acanthomorpha group of spiny-rayed fishes (Rosen, 1973), many of which show dramatic changes in the size, shape, or number of axial skeletal structures. A key evolutionary innovation of this group is the development of stiff, unsegmented bony spines anterior to the median dorsal and anal fins. These dorsal spines can be raised to protect against predators or lowered to facilitate swimming (Wainwright and Longo, 2017). The number, length, and morphology of bony spines differ substantially among species, and the spines can be freestanding or incorporated into segmented rays within the dorsal and anal fins (Mabee et al., 2002). Recent studies have begun to reveal how spines form and grow within the median fin fold of developing fish (Höch et al., 2021; Howes et al., 2017; Roberts Kingman et al., 2021a). However, little is known about the detailed molecular changes that underlie the diverse patterns of spines seen in different fish species.

Sticklebacks form a diverse clade of fish within the Acanthomorpha. Multiple genera of sticklebacks live in the marine and freshwater environments around the northern hemisphere and diverged over 16 million years ago (Aldenhoven et al., 2010; Kawahara et al., 2009; Mattern, 2006). The most well studied of these species, *Gasterosteus aculeatus*, also known as the threespine stickleback, colonized many new freshwater postglacial habitats from the oceans following widespread melting of glaciers at the end of the last ice age, approximately 12,000 years ago (Bell and Foster, 1994). In new freshwater environments containing different food sources and predators, *Gasterosteus* populations have evolved substantial differences in craniofacial structures, vertebrae, and the number of defensive bony plates and spines along the A-P body axis (Bell and Foster, 1994). Many of the recently evolved populations show loss or reduction of structures previously present in ancestral forms, including loss of armor plates, loss of the pelvic hind fins, reduction of spine lengths, and reduction of body pigmentation (Chan et al., 2010; Colosimo et al., 2005; Howes et al., 2017; Miller et al., 2007). However, recently derived populations can also evolve increases in size or number of structures, including increased body size, increased number of teeth, increased spine length, and increased number of spines in the dorsal midline (Cleves et al., 2014; Moodie, 1972; Roberts Kingman et al., 2021a; Spoljaric and Reimchen, 2011).

Here, we use genetic and genomic approaches in two different stickleback genera to study the molecular mechanisms involved in spine patterning changes in natural populations. Our studies provide new evidence to support the long-standing hypothesis that mutations in the *cis*-regulatory regions of *Hox* genes underlie adaptive evolution of skeletal patterns along the A-P body axis of wild vertebrate species.

## Results

### QTL mapping of stickleback spine number and length in *Gasterosteus aculeatus*

To study the genetics of spine number in *Gasterosteus aculeatus*, we generated a large F2 cross by crossing a wild-caught female freshwater stickleback from Boulton Lake, British Columbia, Canada and a wild-caught male marine stickleback from Bodega Bay, California, USA. The Bodega Bay male fish had the three dorsal spines typically seen in *Gasterosteus aculeatus*. The Boulton Lake female fish had two dorsal spines, as is true for 80% of the stickleback fish found in the lake; the other 20% of fish have three dorsal spines (Reimchen, 1980). We intercrossed F1 males and females that each had three dorsal spines and raised 590 F2 offspring from the largest family (Figure 1A). Most F2 individuals had three dorsal spines, but six had two dorsal spines, and twenty-one had four dorsal spines. For comparison of phenotypes among fish that varied in spine patterns, we numbered spines from anterior to posterior, with the posterior-most spine immediately in front of the dorsal fin being called dorsal spine last (DSL). Therefore, a four-spine *Gasterosteus* has dorsal spine 1 (DS1), dorsal spine 2 (DS2), dorsal spine 3 (DS3), and dorsal spine last (DSL), which we refer to as a high-spine phenotype. A typical three-spine *Gasterosteus* has dorsal spine 1 (DS1), dorsal spine 2 (DS2), and dorsal spine last (DSL), which we refer to here as a low-spine phenotype in the context of this study (Figure S1A).

**Figure 1.**
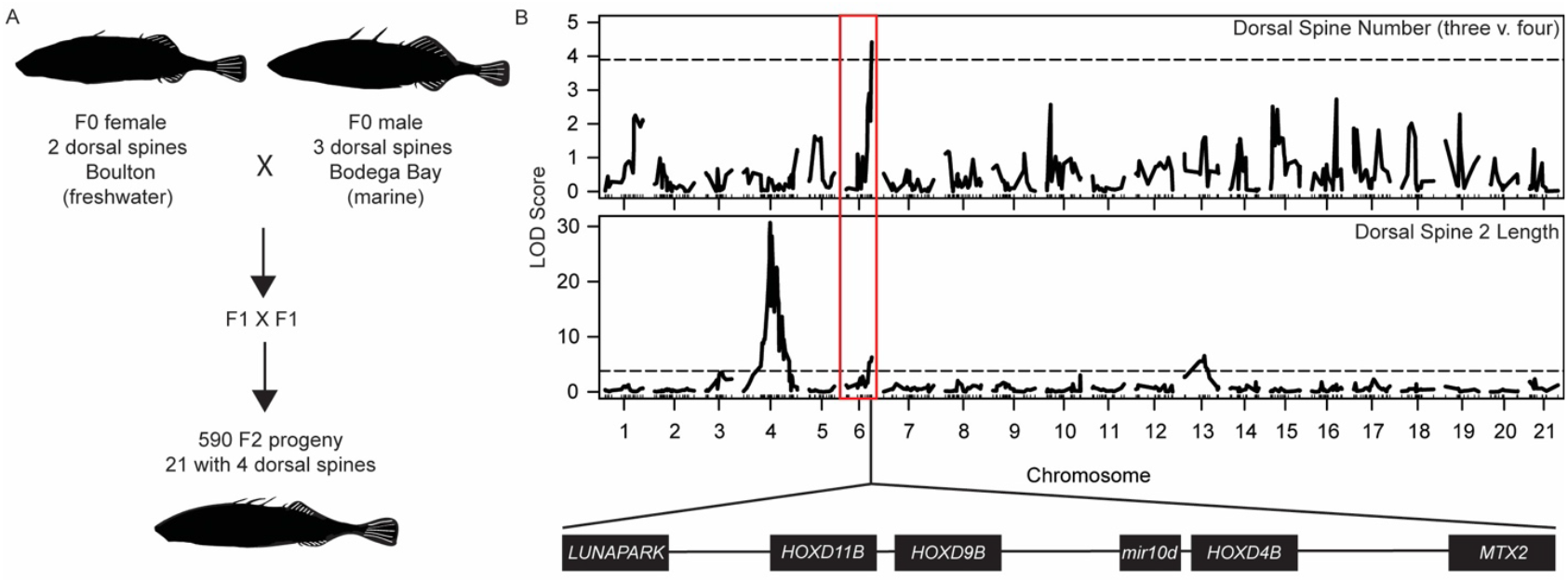
Increased dorsal spine number and decreased dorsal spine 2 length maps to a chromosome 6 region containing the *HOXDB* cluster. **A**. *Gasterosteus* QTL mapping cross. **B**. QTL scan results for dorsal spine number (three-versus four-spine) and DS2 length from F2 progeny of the Boulton Lake and Bodega Bay cross. The x-axis shows the chromosomes in the *Gasterosteus* genome, and the y-axis shows the LOD score for spine number (top) and length of DS2 (bottom). The QTL peak on the distal end of chromosome 6 includes the *HOXDB* cluster drawn below the plot. The major peak for DS2 length on chromosome 4 contains the *EDA*-*MSX2A*-*STC2A* supergene complex, which has been described elsewhere (Howes et al., 2017; Roberts Kingman et al., 2021a). Dashed lines: genome-wide significance threshold based on permutation testing.

To examine the genetic basis of morphological phenotypes along the A-P body axis, we genotyped 340 fish from the family using a custom SNP array (Jones et al., 2012a) and phenotyped the fish for the number of dorsal spines, the length of dorsal spines, the number of flat bony plates that form in the dorsal midline or at the base of spines (pterygiophores), and the number of abdominal, caudal, and total vertebrae (Figures 1B and S1). There were not enough two-spine fish for the mapping of the two-versus three-spine trait. When mapping three-versus four-spine as a categorical trait, we detected a significant quantitative trait locus (QTL) on the distal end of chromosome 6. The same chromosome region also scored as a significant QTL for DS2 length. The allele linked to both the increase in spine number and the decrease in DS2 length was the allele inherited from the freshwater Boulton parent. None of the vertebral traits mapped to the distal end of chromosome 6 (Figure S1), suggesting that the effect of this chromosome region was specific to patterning dorsal spine number and length but not to axial patterning as a whole. Previous studies of other stickleback populations have identified other loci that control vertebral number (Berner et al., 2014; Miller et al., 2014).

### *HOXDB* is in the candidate interval and expressed in *Gasterosteus* spines

The distal end of chromosome 6 contains the *HOXDB* locus in *Gasterosteus*. While this locus is unannotated in the *Gasterosteus aculeatus* reference genome (gasAcu1, (Jones et al., 2012b)), previous studies of *Hox* clusters across multiple species suggest the locus includes three genes (*HOXD11B, HOXD9B*, and *HOXD4B*) and one microRNA (*miR-10d*) (Hoegg et al., 2007). *Hox* genes are known to be expressed in the neural tube and somites as the body axis forms early in development (Ahn and Gibson, 1999b, 1999a). To investigate *HOXDB* gene expression in sticklebacks, we used *in situ* hybridization during embryonic axis formation (stage 19/20; Swarup, 1958). *HOXD4B* was expressed in the hindbrain, neural tube, and anterior-most somites, *HOXD9B* was expressed more posteriorly in the somites and neural tube, and *HOXD11B* was expressed in the most posterior somites and tailbud (Figure S2), consistent with similar colinear patterns found in many other organisms (Burke et al., 1995; Carroll et al., 2005).

Dorsal spines form weeks after early embryonic patterning within a median fin that encircles the developing stickleback (stages 28-31; Swarup, 1958). To examine expression at later stages, we designed a knock-in strategy to introduce an eGFP reporter gene upstream of the endogenous *HOXD11B* locus using CRISPR-Cas9 (Figure 2A). This was done to assess more time points than possible by *in situ* hybridization and also because probe penetration became a problem at later time points. The reporter line was generated in an anadromous *Gasterosteus* background from the Little Campbell River, British Columbia, Canada, a typical three-spine *Gasterosteus* stickleback population that migrates between marine and freshwater environments (Hagen, 1967). At stage 19/20, we saw GFP expression in the posterior somites and tail bud, a pattern that recapitulated the *HOXD11B in situ* hybridization expression already observed at the same early embryonic stage (Figure 2B, Figure S2C). Later in development at stage 31, when the dorsal spines are forming, we saw expression in the posterior half of the fish (Figure 2C) and in the dorsal fin fold between the DS2 and DSL, the DSL, and the dorsal fin (Figure 2D). We also saw expression in the anal fin and anal spine. This reporter expression suggests that *HOXD11B* is expressed both in early development and later during dorsal spine formation in the median fin (a conclusion also supported by RNA-sequencing experiments, see below and Figure S4).

**Figure 2.**
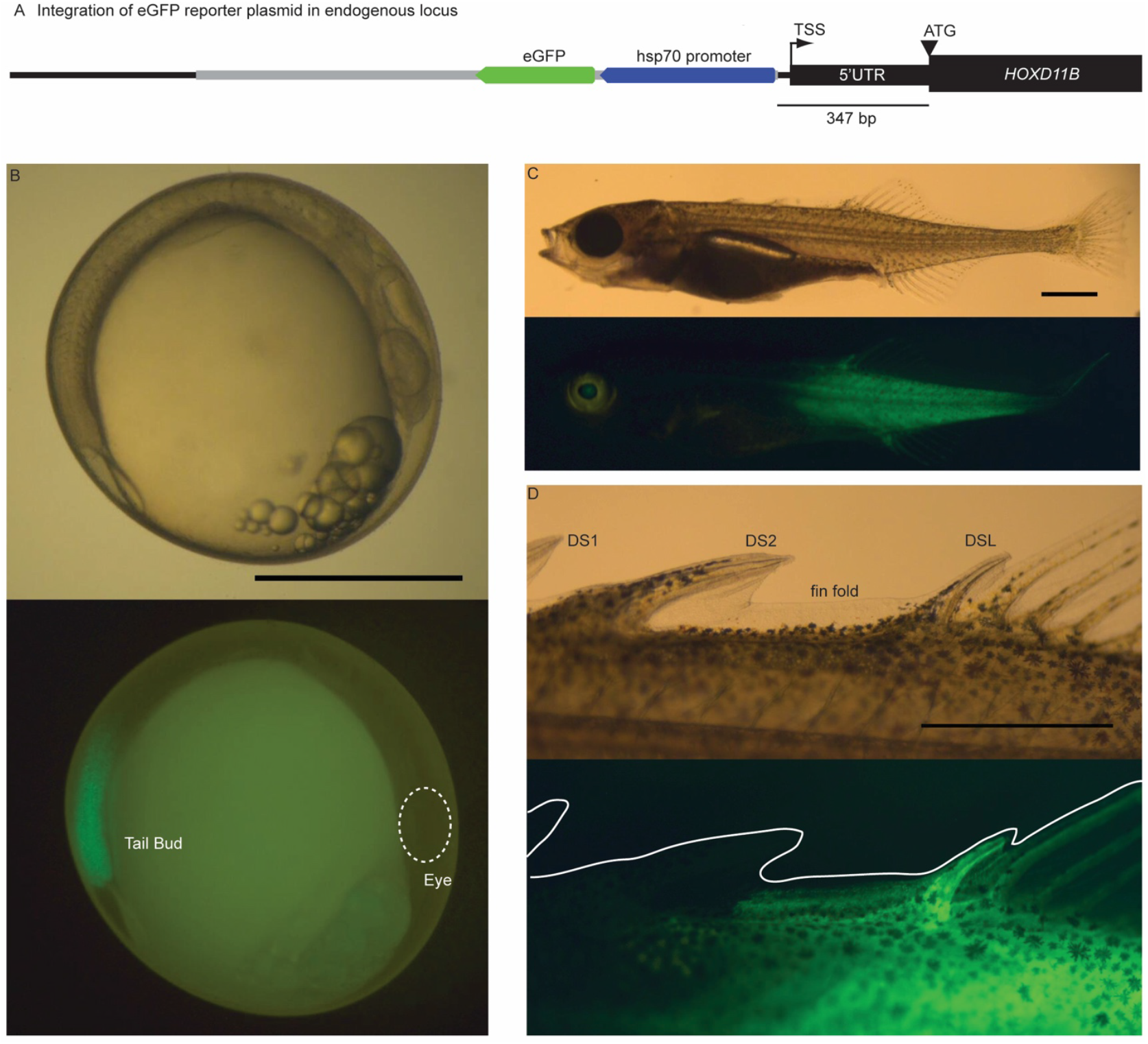
GFP reporter upstream of *Gasterosteus HOXD11B* shows expression in somites, fins, and spines. **A**. Schematic of the integration of the reporter plasmid by CRISPR-Cas9 upstream of the endogenous *HOXD11B* locus of low-spine *Gasterosteus*. The plasmid is in gray; eGFP is in green; hsp70 promoter is in blue; the endogenous locus is in black. **B**. Embryonic expression in the somites in the tailbud at Swarup stage 19/20. The pattern recapitulates the *in situ* hybridization results for *HOXD11B* (Figure S2C). The dotted circle shows the location of the eye. **C**. GFP expression at Swarup stage 31 when the dorsal spines are formed. GFP expression was seen in the posterior half of the fish. **D**. In the dorsal structures, GFP expression was seen in the fin fold between DS2 and DSL, the DSL, and the dorsal fin. All scale bars are 1 mm.

To determine if *HOXDB* genes are functionally important for dorsal spine patterning, we used CRISPR-Cas9 to target the coding region of *HOXD11B* in typical anadromous low-spine *Gasterosteus* (Little Campbell, British Columbia, Canada). Fish in the F0 generation that were mosaic for different mutations in the coding region of *HOXD11B* showed significantly longer DSL compared to their uninjected control siblings (Figure 3, two-tailed t-test; p=7E-06, n=18 control and 23 injected). The anal spine (AS) was also significantly longer in the F0 injected fish (Figure 3, two-tailed t-test; p=0.006, n=18 control and 23 injected). In addition to the spine length, we also saw an effect on the number of bony basal plates or pterygiophores along on the dorsal midline of the fish (Figure S1). While low-spine *Gasterosteus* develop with either one or two blank (non-spine bearing) pterygiophores between DS2 and DSL, all CRISPR-Cas9 targeted fish developed with only one pterygiophore (n=5/18 control with two blank pterygiophores; n=0/23 injected F0 mutants with two blank pterygiophores, two-tailed Fisher’s exact test p=0.01). To further validate the CRISPR results, we also tested the effect of *HOXD11B* targeting in a second anadromous population (Rabbit Slough, Alaska, USA). Again, we observed a significant effect on the length of both the DSL and AS (Figure S3, two-tailed t-test; DSL p=1E-13, AS p=4E-08, n= 38 injected and n=30 control from 3 clutches combined). Because the Rabbit Slough control population does not have variable pterygiophore numbers, an effect on blank pterygiophores could not be examined. These results show that *HOXD11B* is functionally important to the patterning of the dorsal spines, specifically DSL length and pterygiophore number.

**Figure 3.**
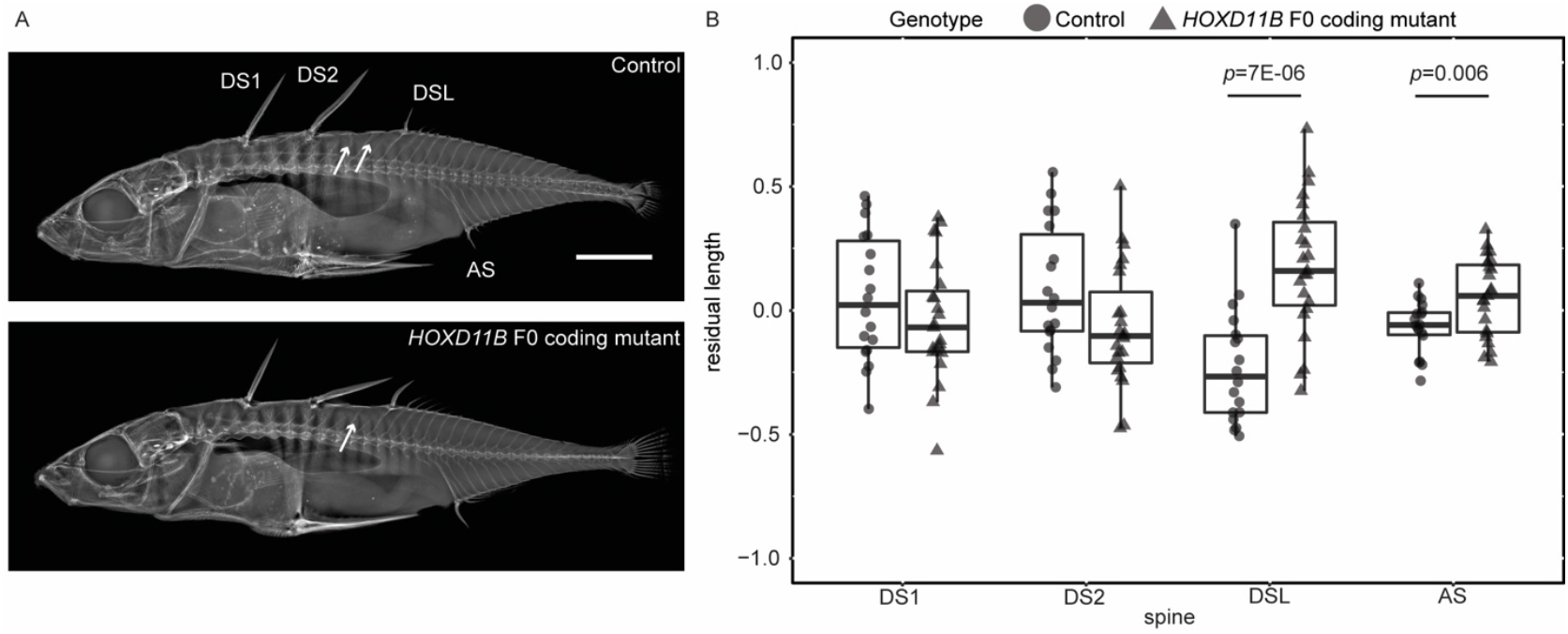
Coding mutations in *HOXD11B* change *Gasterosteus* spine length. **A**. X-ray of an uninjected *Gasterosteus* (top) and *Gasterosteus* injected at the single-cell stage with Cas9 and an sgRNA targeting the coding region of *HOXD11B* (bottom). DS1, dorsal spine 1; DS2, dorsal spine 2; DSL, dorsal spine last; AS, anal spine. Arrows point to blank pterygiophores between DS2 and DSL; two blank pterygiophores are only seen in uninjected fish (see text). The scale bar is 5 mm. **B**. Quantification of length differences in dorsal and anal spines. The y-axis is the residual after accounting for the fish standard length (Figure S1A). DSL and AS were significantly longer in the injected *HOXD11B* F0 coding mutants compared to the controls (two-tailed t-test, n= 18 control and n= 23 injected, DSL p=7E-06, AS p=0.006). There was no significant difference in spine length in DS1 and DS2.

### *HOXDB* gene expression is expanded in dorsal spines of high-spine *Gasterosteus*

To examine whether four-spine/high-spine *Gasterosteus* fish have *cis*-acting regulatory changes in *HOXDB* gene expression, we generated F1 hybrids between low-spine and high-spine stocks and used RNA-sequencing to look for allele-specific expression patterns that were detectable even when both alleles were present in the same trans-acting environment. The hybrids were generated by crossing Little Campbell River anadromous fish, which predominantly have three dorsal spines (referred to as low-spine), with a stock descended from the QTL progeny that carry the Boulton *HOXDB* allele and predominantly show four or five spines (referred to as high-spine, see methods, RNA-sequencing section). In this cross, 77% of the 57 F1 hybrids had three dorsal spines, 21% had four dorsal spines, and one fish had five dorsal spines. RNA was isolated from micro-dissections of each dorsal spine (DS1, DS2, DS3 (if present), DSL), blank pterygiophore (Pter), dorsal fin (DF), and anal fin (AF) at the developing fin fold stage (displayed in Figure 4A). RNA was also isolated from whole embryos at embryonic stage 19/20.

**Figure 4.**
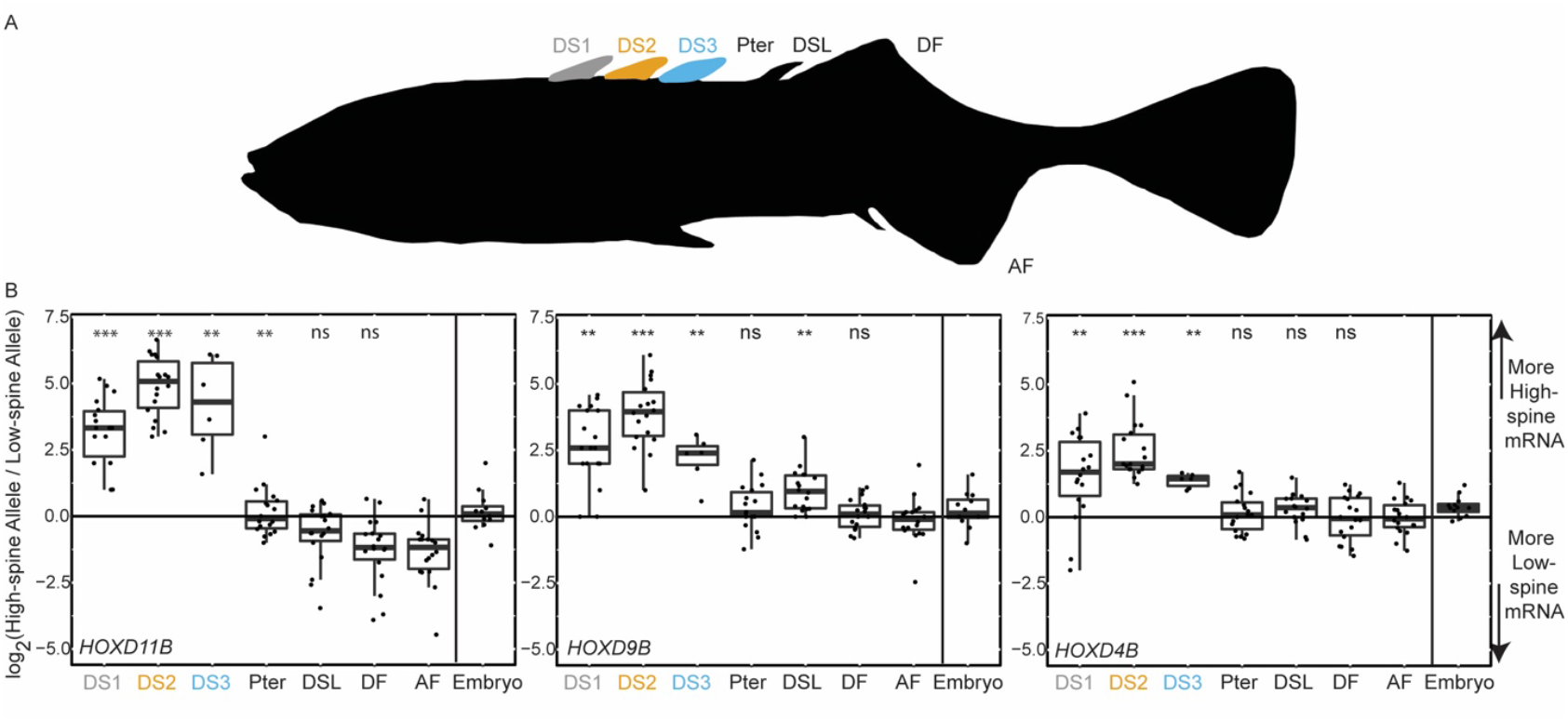
*HOXDB* genes show *cis*-acting expression differences in *Gasterosteus* spines. **A**. F1 progeny were generated in a cross between a low-spine *Gasterosteus* and a high-spine *Gasterosteus*, and tissues were isolated from up to seven indicated locations (DS1, DS2, DS3 (if present), Pter, DSL, DF, AF) to measure allele-specific gene expression in the fin fold stage. Note DS3 only developed in some F1 progeny, so this location has fewer samples (n=6 for DS3; n=18 for all other tissues). **B**. The box plots show ratios of high- to low-spine allele expression at each of three *HOXDB* genes. The y-axis is the log2 of the high-spine versus low-spine read ratio at a SNV (black line: equal expression at log2 ratio of 0). The x-axis shows the seven tissues collected from fin-fold stage fish arranged from anterior to posterior, as well as the sample collected from earlier whole embryos (Embryo). SNVs scored for each dorsal tissue compared to the anal fin: *HOXD11B*, chrVI:17756571; *HOXD9B*, chrVI:17764664; and *HOXD4B*, chrVI:17783616 (*gasAcu*1-4). *** indicates that p ≤ 1E-6 and ** indicates p ≤ 1E-3 by Mann Whitney U test. All alleles with 0 reads have been replaced with 0.5 for graphical representation purposes and statistical analysis.

All three *HOXDB* genes were expressed in the whole embryo samples from stages 19/20 (Figure 4B). Reads from RNA sequencing were assigned to low- or high-spine *HOXDB* alleles using exonic single nucleotide variants (SNVs) that differ between Little Campbell or Boulton haplotypes. *HOXD9B* showed no significant allele-specific expression differences at 5 different informative SNVs. *HOXD11B* showed differences at three out of eight SNVs (binomial test p< 0.01), and *HOXD4B* showed differences at all three of the informative SNVs (binomial test p< 0.001) (Figure 4B). Different results for different SNPs may reflect the heterogeneity of expression locations and gene isoforms present in whole embryos. Overall, there were no striking differences in expression between the two alleles at the embryonic timepoint.

At the later fin fold stage, we sequenced dissected tissues from twelve three-spined and six four-spined F1s. We compared the expression in the dorsal spines and fins to anal fin expression as a control. DS1, DS2, and DS3 showed allele-specific expression differences of all three *HOXDB* genes. In each case, substantially higher expression was seen from the high-spine parent allele (Figure 4B). The expression differences were seen in all F1 hybrid siblings, regardless of whether they had a three- or four-spined phenotype. Elevated expression of the high-spine allele was not seen for the Pter, DSL, or DF locations (Figure 4B). In DS1 and DS2, almost all detectable sequence reads for all three *HOXDB* genes came from the high-spine *Gasterosteus* allele, and very few or none came from the low-spine *Gasterosteus* allele. This is consistent with the previous patterns observed with the *HOXD11B* low-spine GFP reporter line, which showed expression of the low-spine allele at posterior, but not anterior, locations in the fin fold (Figure 2D). The elevated expression coming from the high-spine allele led to a significant positive log2 ratio of high-spine to low-spine expression in each of the dorsal spines when compared to the anal fin (DS1: *HOXD11B* (chrVI:17756571) p= 3E-07; *HOXD9B* (chrVI:17764664) p= 6E-06; *HOXD4B* (chrVI:17783616) p= 1E-05; DS2: *HOXD11B* (chrVI:17756571) p= 3E-07; *HOXD9B* (chrVI:17764664) p= 4E-07; *HOXD4B* (chrVI:17783616) p= 4E-07; DS3: *HOXD11B* (chrVI:17756571) p= 4E-04; *HOXD9B* (chrVI:17764664) p= 8E-04; *HOXD4B* (chrVI:17783616) p= 6E-04, all p-values by Mann Whitney U test). Similar results were seen for all SNVs that were scoreable in the three *HOXDB* genes (*HOXD11B* 9 SNVs; *HOXD9B* 4 SNVs; *HOXD4B* 2 SNVs).

### *HOXDB* is associated with dorsal spine number and length in *Apeltes quadracus*

To determine if other stickleback genera use the same locus to control dorsal spine patterning, we conducted an association mapping study in *Apeltes quadracus*. As their scientific species name suggests, *Apeltes “quadracus”* typically has four dorsal spines. However, multiple populations in the maritime provinces of Canada have previously been identified that show a high incidence of either low- or high-spine fish (Blouw and Hagen, 1984a) (see Figures 5 and S5 for further details of anatomy). The *Apeltes* spine number differences are heritable and correlated with ecological conditions across different geographic locations (Blouw, 1982; Hagen and Blouw, 1983). We sampled from two populations in Nova Scotia, one (Louisbourg Fortress) with predominantly five dorsal spines (range from three to six, Figure 5A), and one (Tidnish River 3) with predominantly four dorsal spines (range from two to six). Approximately equal numbers of low-spine (two to four spines) and high-spine (five to six spines) individuals were genotyped across the *HOXDB* locus (Louisbourg Fortress n= 211 total, 1 three-spine, 104 four-spine, 99 five-spine, 7 six-spine; Tidnish River n=121 total, 1 two-spine, 1 three-spine, 59 four-spine, 59 five-spine, 1 six-spine). A highly significant association was seen between spine number in wild fish and the genotypes at two markers located between *HOXD9B* and *HOXD11B* (Figure 5B, Black line). At the peak marker (*AQ-HOXDB*_6), fish homozygous for the AA allele had an average of 5.1 spines (SD = 0.4) while fish homozygous for the GG allele had an average of 4.2 spines (SD=0.5).

**Figure 5.**
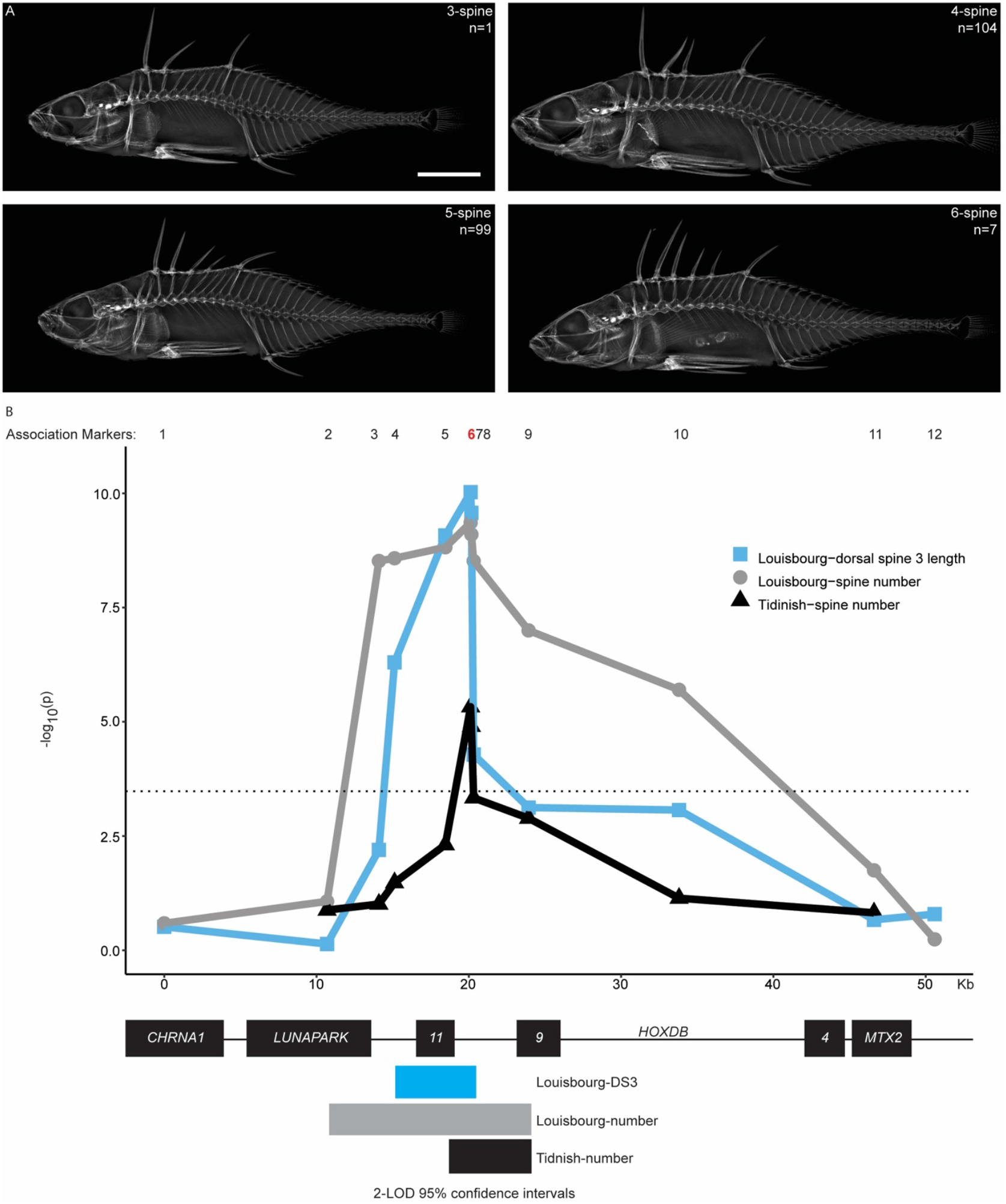
Spine number and dorsal spine 3 (DS3) length are associated with the *HOXDB* locus in *Apeltes quadracus*. **A**. X-rays of *Apeltes quadracus* from Louisbourg Fortress with three to six dorsal spines. Scale bar is 5 mm. **B**. Association mapping of *Apeltes quadracus* from Louisbourg Fortress (n=211; n=1 three-spine, n=104 four-spine, n=99 five-spine, n=7 six-spine) and Tidnish River 3 (n=121; n=1 six-spine, n=59 five-spine, n=59 four-spine, n=1 three-spine, n=1 two-spine). Both populations show a significant association between spine number and the *HOXDB* locus. *Apeltes quadracus* from Louisbourg Fortress were also phenotyped for dorsal spine length and show a significant association between DS3 length and the *HOXDB* cluster. The markers used are displayed across the top (1-12), and the peak marker (6) is highlighted in red. The dotted line represents the Bonferroni corrected significance threshold at α = 0.05. The 95% confidence intervals (2-LOD) for spine number are denoted by the bars on the bottom (gray for Louisbourg, black for Tidnish). The overlapping 95% confidence interval for spine number is ~5750 bp from the third exon of *HOXD11B* to the first exon of *HOXD9B*. The smallest interval shared by both spine number and spine length intervals is ~2kb including *HOXD11B* exon 3 and part of the intergenic region between *HOXD9B* and *HOXD11B*. Additional anatomical details and association plots for other spine lengths are shown in Figure S5.

To test whether, as in *Gasterosteus*, the *Apeltes HOXDB* cluster is also associated with changes in lengths as well as numbers of spines, we measured the length of the individual dorsal spines and anal spines in the Louisbourg Fortress fish. The spines were numbered similarly to *Gasterosteus*, with a three-spine *Apeltes* having from anterior to posterior DS1, DS2, and DSL, and a six-spine *Apeltes* having DS1, DS2, DS3, DS4, DS5, and DSL (Figure S5A). Comparing spine lengths and genotypes showed the DS3 length was strongly associated with the genotypes in the *HOXDB* region (Figure 5B, blue line), but the other spines lengths were not (Figure S5). The genotype at the peak marker in the *HOXDB* cluster (*AQ-HOXDB*_6) explained 22% of the overall variance in DS3 length of wild-caught fish.

The minimal genomic interval that was shared by both the spine number and spine length associations was approximately ~2kb, including *HOXD11B* exon 3 and part of the intergenic region between *HOXD9B* and *HOXD11*B (Figure 5B). Based on whole genome DNA sequencing from Louisbourg (n=2) and RNA-sequencing data (n=14) (see methods sections: *Apeltes* genome and assembly and RNA-sequencing), no sequence variation was found in the protein-coding regions of *HOXD11B* or *HOXD9B*. The peak marker for both spine number and length associations was a change of two adjacent base pairs from GG to AA in the intergenic region. Together, these results suggest that the increased spine number and increased DS3 length in some *Apeltes* likely arise from a regulatory difference that maps in the non-coding interval between *HOXD9B* and *HOXD11B*.

### *Apeltes HOXDB* genes show *cis*-regulatory differences in spine expression

To further test for possible *cis*-acting regulatory differences in *Apeltes HOXDB* genes, we generated F1 hybrids carrying contrasting *Apeltes* haplotypes in the key genomic interval and carried out RNA-sequencing on the spines, blank pterygiophore, and dorsal and anal fins at the fin fold stage (Figure 6A). While there were no sequence differences in the protein coding portions of the *HOXDB* genes, the 3’ UTRs of *HOXD9B* and *HOXD11B* had variants that could be used to determine the expression level coming from the genotypes associated with low-spine (L) or high-spine number (H) in the association study (Figure 6B). In F1 fish carrying one L-haplotype and one H-haplotype, significantly higher expression was seen from the *HOXD9B* and *HOXD11B* genes of the L-haplotype (Figure 6C). The difference was most pronounced and statistically significant in DS3, the same spine whose overall length was associated with genotypes in the *Apeltes HOXDB* region (DS3:*HOXD9B* (chr06:16028519) p= 3E-7; *HOXD11B* (chr06:16020516) p=9E-4 (Fisher’s Exact Test)).

**Figure 6.**
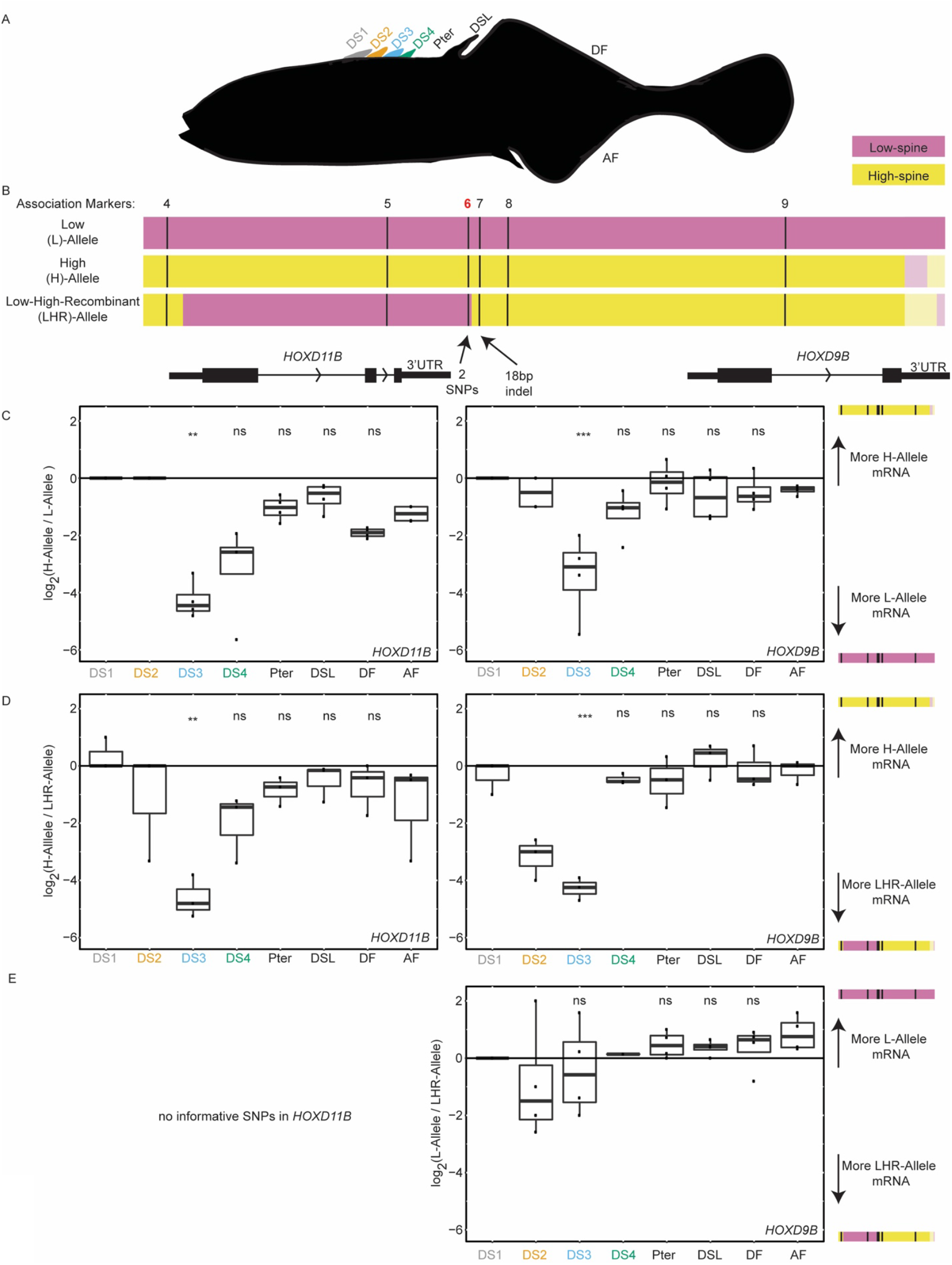
*HOXDB* genes show *cis*-acting expression differences in *Apeltes* spines. **A**. Outline of *Apeltes* fin fold stage fry. Tissues were isolated from up to eight indicated locations (DS1, DS2, DS3, DS4, Pter, DSL, DF, AF) to measure allele-specific gene expression in the fin fold stage. DS4 only developed in some F1 progeny, so this location has fewer samples. **B**. Schematics showing the three *HOXDB* haplotypes that segregate in the allele-specific expression cross. Black lines indicate the position of association mapping markers used to identify the haplotypes. Pink indicates regions where genotypes match those associated with low-spine phenotypes in the association analysis (Figure 5B). Yellow indicates regions where genotypes match those associated with high-spine phenotypes in the association analysis. Lighter shading on the right indicates regions where marker association is not known, but DNA sequence differences are shared between haplotypes of the same color. **C**. Box plots showing the allele-specific expression ratios in all the tissues dissected at fin fold stage from F1 fish heterozygous for H and L haplotypes. Reads from DS3, DS4, Pter, DSL, and DF were compared to reads from AF to determine significance (by Fisher’s Exact Test: *HOXD9B* (chr06: 16028519) p= 3E-7; *HOXD11B* (chr06:16020516) p=9E-4). DS1 and DS2 were not assessed because the read counts were too low (Figure S4). **D**. Box plots showing the allele-specific expression ratios in all the tissues dissected at fin fold stage from fish heterozygous for LHR and H haplotypes. Allele-specific expression was seen in DS3 when compared to the AF for both *HOXD11B* and *HOXD9B* (by Fisher’s Exact Test: *HOXD9B* (chr06: 16027923) p= 9E-8; *HOXD11B* (chr06:16020516) p=1E-3). **E**. Box plot showing the allele-specific expression ratios in all the tissues dissected at fin fold stage from F1 fish heterozygous an L and LHR haplotypes (by Fisher’s Exact Test in DS3: *HOXD9B* (chr06: 16027923) p=0.08). Only *HOXD9B* is shown because there were no informative SNPs in *HOXD11B* between the L and LHR haplotypes. ** p ≤ 1E-3 *** p ≤ 1E-6.

Some of the F1 fish generated for the allele-specific expression experiment carried both an H haplotype and a recombinant haplotype that we termed the low-high-recombinant (LHR) haplotype (Figure 6B). These fish showed an allele-specific expression pattern similar to the one observed in fish heterozygous for an H and L haplotype (Figure 6D), with more expression of *HOXD9B* and *HOXD11B* coming from the LHR haplotype (DS3: *HOXD9B* (chr06:16027923) p= 9E-8; *HOXD11B* (chr06:16020516) p=1E-03 (Fisher’s Exact Test)). In contrast, fish heterozygous for the LHR and L haplotypes showed no significant difference between the *HOXD9B* expression patterns (DS3: *HOXD9B* (chr06:16027923) p=0.08, *HOXD11B* was not scored due to a lack of distinguishing markers between LHR and L) (Figure 6E). Thus, at a gene expression level, the LHR haplotype behaved more like the L haplotype than the H haplotype. Similarly, at the phenotypic level, F1 individuals heterozygous for the L and LHR haplotypes typically had low spine numbers, resembling fish homozygous for the L haplotype, while fish homozygous for the H haplotypes had higher spine numbers (L/L fish: 16/16 with three or four spines; L/LHR fish: 15/18 with three or four spines, 3/18 with five spines; H/H fish: 16/16 with five or six spines). These results suggest the key genomic interval controlling both gene expression differences and phenotypic differences between the L/LHR and H haplotypes maps to the minimal ~5 kb region shared between the L and LHR haplotypes (pink region on the left side of Figure 6B).

### Identification of a spine enhancer and genomic changes in both *Gasterosteus* and *Apeltes*

To search for possible *cis*-regulatory sequences contributing to *HOXDB* expression variation, we looked for conserved non-coding sequences and open chromatin domains located in the minimal interval defined by the association (Figures 1 and 5) and gene expression studies (Figures 4 and 6). This identified one ~500 bp region (Figure 7A) found in both *Apeltes* and *Gasterosteus* that is conserved by phastCons alignment to Tetraodon, Medaka, and Fugu (Siepel et al., 2005). This small conserved region contained the peak scoring marker in the *Apeltes* association study (two adjacent base pairs changed from GG to AA). This conserved non-coding region also corresponds to a region of open chromatin in medaka embryos at stages equivalent to those where we see *HOXDB* expression in sticklebacks by *in situ* hybridization (Marlétaz et al., 2018).

**Figure 7.**
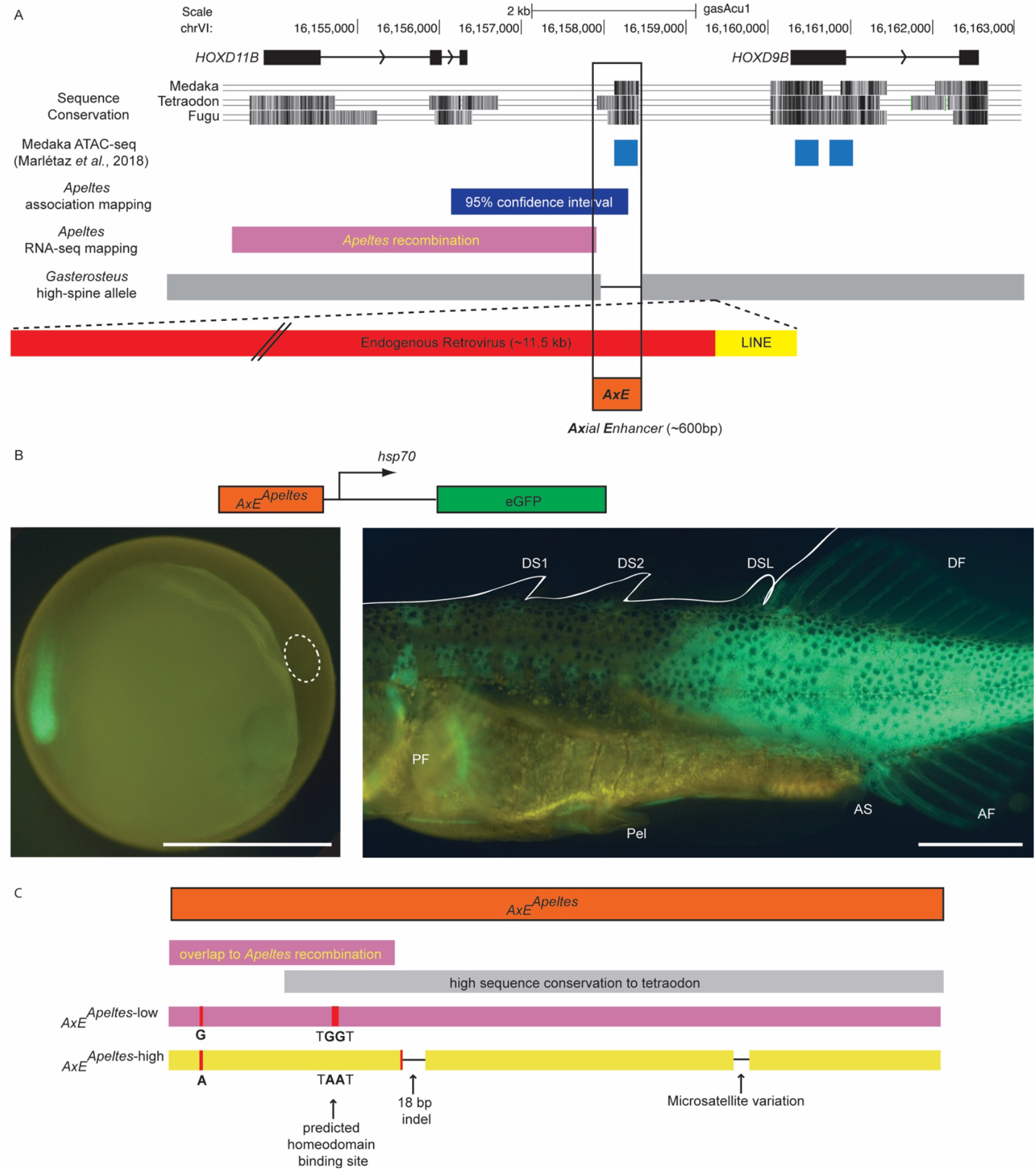
The genomic region between *HOXD11B* and *HOXD9B* contains a conserved axial enhancer showing sequence changes in both *Gasterosteus* and *Apeltes*. **A**. The exons of *HOXD11B* and *HOXD9B* are shown in *Gasterosteus* (gasAcu1) genomic coordinates. Sequence Conservation: phastCons conserved sequence regions identified in exons and in a ~500 bp intergenic region from comparisons between fish genomes. The conserved non-coding region overlaps an ATAC-seq peak from Medaka embryonic stage 19 (Marlétaz et al., 2018) and partially overlaps the genomic intervals defined by spine phenotype and RNA-expression changes in *Apeltes*. In high-spine *Gasterosteus*, the conserved region is deleted (as indicated by a black line between the two gray boxes), and an endogenous retrovirus and LINE sequence are inserted (drawn in red and yellow, respectively). **B**. ~600 bp regions from low-spine and high-spine *Apeltes* were cloned into a *Tol2* GFP expression construct and injected into *Gasterosteus* embryos. Both versions drove expression in the tail bud of embryos (left), and the fin fold, spines, and dorsal, anal, and caudal fins of stage 31 fry (right), confirming the region acts as an enhancer. Scale bar is 1 mm. **C**. There are four sequence differences in the *AxE* region of high- and low-spine *Apeltes* alleles: one microsatellite variation, an 18bp indel, a SNP, and two adjacent SNPs. Only the single and the two adjacent SNPs are within the region implicated by *Apeltes* recombination and RNA-sequencing differences (pink bar).

We cloned the *Apeltes* region from both the L and H haplotypes (611 bp in L; 587 bp in H) and tested whether the sequence could drive GFP reporter gene expression in transgenic enhancer assays. Because *Apeltes* fish have very small clutch sizes, constructs were injected into *Gasterosteus* embryos to obtain sufficient transgenic embryos for analysis. The ~600 bp non-coding constructs both drove expression at embryonic time points in the tail of transgenic embryos in a similar pattern to that seen in the *in situ* hybridizations for *HOXD9B* and *HOXD11B* (Figure 7B, left). At later time points, the conserved non-coding regions drove expression in all of the fins (dorsal, caudal, anal), all of the spines (dorsal spines and pelvic spines), and the posterior portion of the fish (Figure 7B, right). Similar patterns were driven by both the L- and H-type *Apeltes* constructs, though we note that differences in both strength and patterns of expression can be difficult to detect in mosaic transgenic fish resulting from random *Tol2* integration (L 611 bp region: n=22 transgenics with bilateral green eyes; n=6/22 pectoral fin, n=8/22 pelvis, n=9/22 dorsal spines, n=11/22 dorsal fin, n=9/22 anal fin, n=10/22 posterior muscle; H 587 bp region: n=19 with bilateral green eyes; n=2/19 pectoral fin, n=4/19 pelvis, n=9/19 dorsal spines, n=11/19 dorsal fin, n=11/19 anal fin, n=10/19 posterior muscle). Given the consistent expression patterns seen in both tail buds and later axial structures of transgenic fish, we refer to the ~600 bp conserved intergenic sequence as an axial enhancer (*AxE)* of the *HOXDB* locus.

Although *AxE* sequences are conserved between *Apeltes* and typical *Gasterosteus*, we were unable to amplify the *AxE* region from the Boulton high-spine allele identified in the *Gasterosteus* QTL cross. We therefore used PacBio long-read sequencing to identify the intergenic region between *HOXD9B* and *HOXD11B* from the *Gasterosteus* Boulton high-spine allele. The sequenced region shows major structural changes, including a deletion which removes almost all of the *AxE* enhancer, and the presence of two transposable elements not present in the low-spine reference genome from Bear Paw Lake ((Jones et al., 2012b) Alaska, USA) fish: a LINE (L2-5_GA) element and an endogenous retrovirus (ERV1-6_GA-I) (Figure 7A). The LINE element is approximately 1 kb and also appears to be present in some additional stickleback populations in the Pacific Northwest (sequencing data from (Roberts Kingman et al., 2021b)). When the LINE element was detected in other populations, it was not associated with the deletion of the *AxE* sequence as seen in Boulton Lake. The endogenous retrovirus insertion was approximately 11 kb containing open reading frames for an envelope and gag-pol proteins, flanked by ~1 kb long terminal repeats (LTRs). Junction sequences for this retroviral insertion near *AxE* were not found in the sequenced genomes of over 200 other sticklebacks from different populations (Jones et al., 2012b; Roberts Kingman et al., 2021b). The Boulton high-spine allele thus shows both the nearly complete loss of *AxE* and the addition of new sequences in the region.

To determine if loss of the *AxE* sequence alone was sufficient to recapitulate the phenotypic effect of a higher spine number and shorter DS2 length in *Gasterosteus*, we deleted the region in low-spine *Gasterosteus* using CRISPR targeting. In mosaic F0 founder fish, no significant effects on spine number were detected. However, DSL and AS were both significantly longer in the F0 injected mutants compared to their control siblings (two-tailed t-test; DSL p=7E-05, AS p=0.004 n=32 control and n=24 injected) (Figure S7). Both the particular spines affected and the direction of phenotypic effects resembled the phenotypes seen when targeting the *HOXD11B* protein-coding region. These results suggest that the *AxE* region is required for normal spine length patterning in *Gasterosteus*, but that additional sequence changes likely contribute to the spine number phenotypes linked to the region.

## Discussion

Vigorous historical debates have existed about whether mutations in homeotic genes are the likely basis of common morphological changes seen in wild animals. Most laboratory or human-selected mutations in the genes are deleterious. In addition, transposable element insertions are strikingly depleted at *Hox* loci, an effect attributed to the likely deleterious consequences of making substantial regulatory changes in genes essential for development and survival (Lander et al., 2001; Simons et al., 2006). On the other hand, the diversity of *Hox* cluster number, composition, and expression patterns, and the powerful effects of *Hox* genes on many phenotypes in laboratory models, have made the genes often-cited candidates for the possible molecular basis of obvious phenotypic differences between wild species, including in sticklebacks (Ahn and Gibson, 1999b, 1999a). *Cis*-acting regulatory differences at *Hox* loci clearly underlie evolutionary differences in trichome and pigmentation patterns in insects, but the underlying molecular changes are still not known (Stern, 1998; Tian et al., 2019). Our studies show that independent regulatory changes have occurred in the *HOXDB* locus of *Gasterosteus* and *Apeltes*, providing a compelling example of *cis*-acting variation in *Hox* genes linked to the evolution of novel adaptive axial skeletal patterns in wild vertebrate species.

### Adaptive significance of dorsal spine number and length

Dorsal spines in sticklebacks play an important role in predator defense. Long spines increase the effective cross-sectional diameter of sticklebacks (Reimchen, 1983) and can provide a survival advantage against gape-limited predators (Hoogland et al., 1956). Prominent spines also provide holdfasts for grappling insect predators and may therefore increase the risk of predation by macroinvertebrates (Marchinko, 2009; Reimchen, 1980). Different predation regimes may thus favor either increased or decreased spine lengths and numbers. The intensity of bird, fish, and insect predation varies across locations, years, and seasons, contributing to a range of different spine phenotypes in natural stickleback populations (Reimchen and Nosil, 2002; Reimchen et al., 2013).

In *Gasterosteus*, fish with four dorsal spines are found at high frequency in a few populations in Alaska and Massachusetts (Bell and Baumgartner, 1984; Bell et al., 1985), though ecological factors acting in these populations are not well characterized. In contrast, Boulton Lake is an extensively studied population where fish typically show two or three dorsal spines, as well as a high incidence of pelvic spine loss (Reimchen, 1980). Detailed seasonal and longitudinal surveys have shown that lower spine numbers in Boulton fish are correlated with a higher intensity of insect predation and higher spine numbers with a higher intensity of bird predation (Reimchen and Nosil, 2002, 2004). Because fish with four dorsal spines have not been seen in the samples of over 20,000 wild-caught Boulton Lake fish, the occurrence of four-spine sticklebacks in the Boulton Lake x Bodega Bay F2 laboratory cross appears to be a transgressive phenotype (Rieseberg et al., 1999) that emerges when Boulton alleles are inherited on a mixed genetic background. We note, however, that the Boulton *HOXDB* region is also linked to shortening of DS2 in the QTL cross. Boulton Lake fish have shorter dorsal spines than marine fish, and we hypothesize that the *HOXDB* allele likely evolved for its contributions to reduced DS2 length in the wild lake population.

*Apeltes quadracus* sticklebacks are named for their typical development of four prominent dorsal spines. However, many wild “*quadracus*” populations in Canada have predominantly three- or five-spines (Blouw, 1982; Blouw and Hagen, 1984a). In an extensive comparison of *Apeltes* spine numbers and environmental variables over 570 different locations, Blouw and Hagen found that increased spine number was correlated with the presence of predatory fish, while decreased spine number was correlated with both more and denser types of vegetation (Blouw and Hagen, 1984b, 1984c). Spine numbers in both *Apeltes* and *Pungitius* (ninespine stickleback) trend in the same direction when both stickleback species are present in the same lake, suggesting that the changes in spine number are selected in response to shared environmental factors, rather than varying randomly (Blouw and Hagen, 1984d). To further study the possible adaptive value of spine number differences, Blouw and Hagan exposed mixed populations of four- and five-spine *Apeltes* to predatory fish and measured differential survival of spine morphs when half of the sticklebacks had been eaten (Blouw and Hagen, 1984c). When vegetation was present, predation was nonselective; however, when vegetation was absent, five-spine fish were less likely to be eaten by perch and trout. Consistent with both the experimental predation experiments and known ecological correlations, stomachs of wild-caught trout contain more four-than five-spine fish, while stomachs of great blue herons contained more five-than four-spine fish (Blouw and Hagen, 1984c). Dorsal spines in sticklebacks thus provide an excellent example of a prominent adaptive structure in vertebrates which evolves in response to different predation regimes in natural environments and diversifies in part through repeated regulatory changes in *Hox* genes.

### *Hox* genes and dorsal midline skeletal patterns

*Hox* genes are well known for controlling the identity of structures in repeating series, including body segments in insects, somite fates in vertebrates, digit identities in limbs, and rhombomere segments in the hindbrain (Carroll et al., 2005). Dorsal spines and pterygiophores represent an additional series of repeating structures found in fish, and many of the spine changes we see in sticklebacks are consistent with identity transformations in the dorsal midline. Prior work has shown that *Hox* phenotypes are often governed by the rule of posterior prevalence, where the posterior-most, highest-numbered *Hox* gene that is expressed in a given region generally controls the fate of that region (Durston, 2012). Therefore, when posterior genes expand in expression, the regions with expanded expression generally acquire a more posterior fate. Conversely, when activity of a *Hox* gene is lost from a region, that region typically acquires a more anterior fate. Because most vertebrates have multiple *Hox* clusters, axial phenotypes are frequently only seen when multiple or all members of a *Hox* paralogous group (PG) are mutated. For example, when all members of the mouse *Hox* PG9 (*Hoxa9, Hoxb9, Hoxc9, Hoxd9*) are mutated, some of the normally rib-less lumbar vertebrae develop ribs like thoracic vertebrae (McIntyre et al., 2007). These vertebrae have thus undergone a homeotic transformation and acquired a more anterior fate.

Our RNA sequencing studies show that many different *Hox* genes in both *Gasterosteus* and *Apeltes* are expressed in the dorsal spines or fins of developing sticklebacks, with the exception of *Hox* PG1, some *Hox* PG13 genes, and all *HOXBB* cluster genes (Figure S4). As in other systems, lower-numbered *Hox* genes tend to be expressed at higher levels at more anterior locations, and higher-numbered genes tend to be expressed at higher levels at more posterior locations. Several *Hox* genes show strong differential expression across different spines and pterygiophores (including *HOXDB* genes), suggesting morphological fates in the dorsal midline are likely influenced by the combined expression of multiple genes.

The *Gasterosteus* high-spine allele with expanded expression of all three *HOXDB* genes is associated with increased spine number and decreased DS2 length. The expanded expression of *HOXD11B* would be predicted to result in a posteriorization of the regions gaining expression. The blank pterygiophore acquires a spine bearing fate, consistent with a shift to a more posterior identity like DSL. The DS2, which is normally the longest, is shortened and thus becomes more like the last dorsal spine, which is the shortest spine. Conversely, knocking down *HOXD11B* expression by CRISPR-Cas9 targeting would be predicted to result in anteriorization of structures, and the increased length of the DSL that we observe is consistent with a shift of DSL to a more anterior and therefore longer spine fate.

The *Apeltes* H-allele that shows reduced *HOXD9B* and *HOXD11B* gene expression in DS3 is associated with both increased spine number and a longer DS3. The number and length phenotypes can both be interpreted as transformations to a more anterior fate. In this model, the appearance of a fifth spine on a normally blank pterygiophore could be explained by partial transformation to a more anterior, spine-bearing fate (analogous to the appearance of thoracic ribs on anteriorized lumbar vertebrae in previous mouse experiments). Anterior spines are normally longer than posterior spines in *Apeltes*, so the increased length of DS3 is also consistent with an anterior transformation.

### Independent sequence changes in *HOXDB cis*-regulatory elements

Our allele-specific expression experiments in F1 hybrids show that the changes in *HOXDB* expression we see in sticklebacks are due to *cis*-acting regulatory differences linked to the *Hox* genes themselves, rather than the secondary consequence of changes in unknown *trans*-regulatory factors. The mapping, association, and transgenic experiments have also identified a particular *cis*-acting enhancer region located between *HOXD9B* and *HOXD11B* that can recapitulate axial expression patterns and shows independent sequence changes in *Gasterosteus* and *Apeltes* with different spine numbers. In *Apeltes*, the most likely sequence difference that mediates changes of *HOXD9B* and *HOXD11B* expression are two adjacent SNPs (marker: *AQ*-6) in *AxE*. These SNPs convert a T<u>GG</u>T sequence in the L-allele to T<u>AA</u>T in the H-allele. They represent the peak marker scored by association mapping and also map within the minimal *cis*-acting recombination interval that controls H vs. L and LHR expression differences when the contrasting alleles are scored in F1 hybrids. Both the T<u>AA</u>T change and a nearby 18 bp indel, which represents the second highest scoring marker (*AQ*-7) in the association mapping, are found in the high-spine fish of two populations on opposite coasts (east and west) of Nova Scotia. Repeated evolution of different five-spine *Apeltes* populations thus likely takes place through a shared underlying molecular haplotype at the *Hox* locus, rather than independent mutations in these different populations. A similar process of allele sharing underlies recurrent evolution of a variety of other phenotypic traits in both sticklebacks and other organisms (Barrett and Schluter, 2008; Colosimo et al., 2005; Martin and Orgogozo, 2013). We note that the derived T<u>AA</u>T sequence in the *Apeltes* five-spine allele creates a predicted core binding motif for a homeodomain protein. Previous studies in *Drosophila* and other organisms have shown that *Hox* genes can autoregulate in positive and negative feedback loops (Bienz and Tremml, 1988; Delker et al., 2019; Irvine et al., 1993). We hypothesize that the creation of a new putative homeodomain binding site located between the *HOXD9B* and *HOXD11B* genes may contribute to the decreased *HOXDB* expression observed with the H-allele. We note, however, that we have not been able to recapitulate the altered expression patterns using *AxE* transgenic reporter constructs integrated at random locations in the genome. Because endogenous *Hox* expression patterns are likely controlled by interactions between multiple long-distance control elements and surrounding topological domains (Montavon et al., 2011; Spitz et al., 2003), the most accurate functional tests of the phenotypic effects of particular mutations will come from scoring those changes at their correct position in the genome. Future advances in genome editing may eventually make it possible to recreate or revert the T<u>GG</u>T and T<u>AA</u>T sequence change at the endogenous *HOXDB* locus in sticklebacks and to further test whether these two adjacent base pair changes are sufficient to alter *Hox* gene expression and spine length or number.

In *Gasterosteus*, the *AxE* enhancer is deleted from the *HOXDB* high-spine allele from Boulton Lake and has been replaced with two transposable elements, one ERV and one LINE. Removing the endogenous *AxE* enhancer by CRISPR targeting does not lead to spine number and DS2 phenotypes, but does recapitulate the DSL length changes seen by targeting *HOXD11B* coding region. The long terminal repeats found in endogenous retroviruses can act as enhancers (Thompson et al., 2016), and we hypothesize that the additional inserted sequences in the Boulton allele underlie broader expression in the dorsal spines and the other phenotypic consequences of the Boulton high-spine allele.

### Repeated use of regulatory changes in morphological evolution

A long-standing question in evolutionary biology is whether the same genetic mechanisms are used repeatedly to evolve similar traits in different populations and species. Although *Gasterosteus* and *Apeltes* last shared a common ancestor over 16 million years ago (Kawahara et al., 2009), our data show that both stickleback groups have made independent *cis*-regulatory changes in the *HOXDB* region which are linked to new dorsal spine patterns in recently evolved, post-glacial populations. The types of mutations made in the *AxE* regulatory region are clearly distinct, and the naturally occurring *Gasterosteus* and *Apeltes* H-alleles lead to contrasting increases and decreases of *HOXDB* expression. Interestingly, the *HOXD* locus also appears to be used repeatedly during horn evolution in mammals. The *HOXD* region shows accelerated evolution and insertion of a novel retroviral element in the diverse clade of species with headgear (horns, antlers, and ossicones, (Wang et al., 2019)). In addition, rare polycerate (four-horned) variants of sheep and goats have recently been shown to have independent mutations in the *HOXD* locus, ranging from a four base pair mutation that alters splicing to a large deletion that removes more than 500 kb of sequence and is lethal when homozygous (Allais-Bonnet et al., 2021; Greyvenstein et al., 2016; Ren et al., 2016). The fish and mammalian results support a growing body of literature that has found repeated use of the same loci underlying similar traits, even though the direction of effect of gene expression and mutational mechanism are often different (Martin and Orgogozo, 2013).

While both of our examples of spine variation in recently diverged populations of *Gasterosteus* and *Apeltes* involve *cis*-regulatory changes, *Hox* coding region mutations may also contribute to diversification of spine patterns over a wider phylogenetic scale. For example, the *Gasterosteidae* family can be separated into five different genera of predominantly three-spine, four-spine, five-spine, nine-spine, and fifteen-spine sticklebacks (*Gasterosteus, Apeltes, Culaea, Pungitius, and Spinachia*, respectively). We note that the coding region of *HOXD11B* shows a high rate of non-synonymous to synonymous substitutions in comparisons across the stickleback family, and the dN/dS ratio is greater than 1.0 for comparisons between *Apeltes* and *Gasterosteus* (Figure S6). This suggests that changes in *HOXD11B* coding regions have likely been under positive selection during the divergence of *Apeltes* and *Gasterosteus*, perhaps contributing to the distinctive patterns of spine length and number that are characteristic of these two different genera.

Spiny-rayed fish are among the most successful of vertebrates, currently making up nearly a third of all extant vertebrate species. The lengths and numbers of spines show remarkable diversity across the Acanthomorpha, including elaborate modifications that have evolved for defense, camouflage, luring prey, or swimming biomechanics (Wainwright and Longo, 2017). Our results show that changes in the dorsal spine patterns of wild fish species have evolved in part through genetic changes in *Hox* genes. Based on the recurrent use of the same *Hox* locus for spine evolution in different stickleback species, we hypothesize that repeated mutations in *Hox* genes may also underlie other interesting changes that have evolved in the axial skeletal patterns of many other wild fish and animal species.

## Acknowledgements

We would like to thank the Semiahmoo First Nation for allowing us to collect sticklebacks from the Little Campbell River (British Columbia, Canada). We thank Brian Summers for setting up the F0 QTL cross, Danielle Desmet for initial measurements of some of the QTL cross fish, Harmony Folse for initial testing of CRISPR GFP-knock-in, Abbey Thompson, Ken Thompson, and Dolph Schluter for help collecting sticklebacks from the Little Campbell River, and Catherine Peichel and Melanie Hiltbrunner for *G. wheatlandi, C. inconstans*, and *S. spinachia HOXD11B* sequences. This work was supported in part by NIH graduate training grant 2T32GM007790 (J.I.W.), pre-doctoral fellowship from the National Science Foundation (T.R.H., G.A.R.K.), a Stanford Graduate Fellowship (G.A.R.K.), a Helen Hay Whitney Postdoctoral Fellowship (A.L.H.), NIH grant R01GM124330 (M.A.B.), and NSERC Discovery Grant RGPIN-2016-04303 (A.C.D.). David Kingsley is an investigator of the Howard Hughes Medical Institute.

## Author Contributions

Conceptualization J.I.W., T.R.H., and D.M.K.; Formal Analysis J.I.W. and T.R.H.; Investigation J.I.W., T.R.H, J.N.A., E.H.A., G.A.R.K., S.D.B., and A.L.H.; Resources T.E.R., M.A.B., C.B.L., A.C.D., and D.M.K; Data Curation J.I.W. and T.R.H.; Writing – Original Draft J.I.W. and D.M.K.; Writing – Review and Editing J.I.W., T.R.H, J.N.A., E.H.A., G.A.R.K., S.D.B., A.L.H., T.E.R., M.A.B., C.B.L., A.C.D., and D.M.K.; Visualization J.I.W.; Supervision D.M.K.; Funding Acquisition C.B.L, A.C.D., and D.M.K

## Supplemental Figures

**Figure S1.**
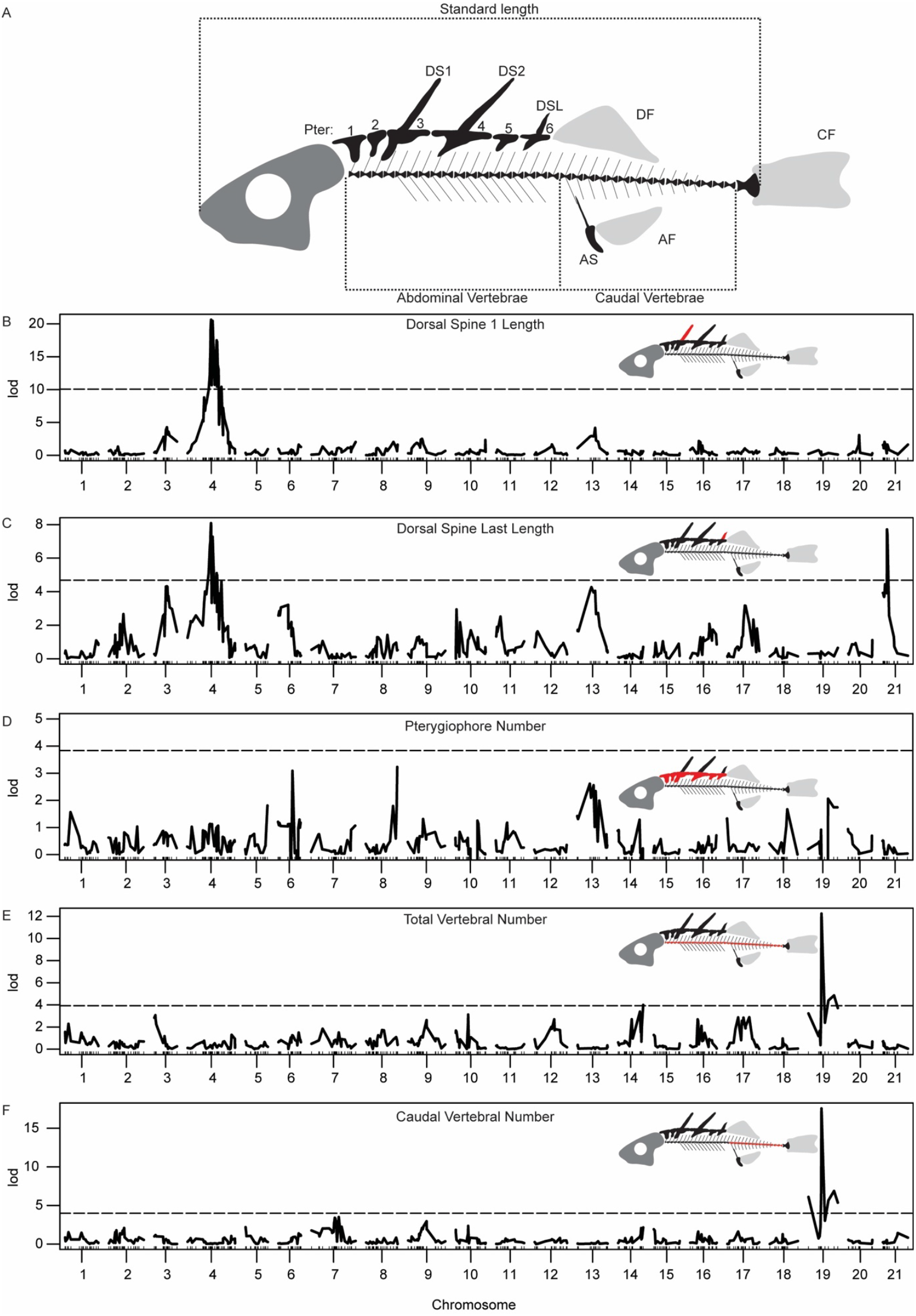
QTL mapping of other spine lengths and axial traits. **A**. Schematic of *Gasterosteus* anatomical features. Most *Gasterosteus* have three dorsal spines that in this study are referred to as dorsal spine 1 (DS1), dorsal spine 2 (DS2), and dorsal spine last (DSL). The dorsal side of the fish also has median bony plates known as pterygiophores, some of which underlie dorsal spines. Typical A-P midline pattern: two non-spine bearing/blank pterygiophores (Pter1 and Pter 2), dorsal spine 1 on pterygiophore 3 (Pter3), dorsal spine 2 on pterygiophore 4 (Pter4), non-spine bearing pterygiophore 5 (Pter5), and dorsal spine last on pterygiophore 6 (Pter6). The three unpaired fins are shown in light gray: dorsal fin (DF), caudal fin (CF), and anal fin (AF). The anal spine (AS) is also indicated on the ventral side of the fish. The standard length shown with the dotted line is from the anterior tip of the jaw to the posterior of the hypural plates. **B**. QTL plot of DS1 length **C**. QTL plot of DSL length **D**. QTL plot of pterygiophore number **E**. QTL plot of total vertebral number **F**. QTL plot of caudal vertebral number. Dotted lines represent genome-wide significance thresholds. Abdominal vertebral number and anal spine length were also tested, but they did not result in any peaks that passed the genome wide significance threshold. The significance threshold (dashed line) is based on LOD scores obtained in 1,000 permutations of the phenotype data (α = 0.05).

**Figure S2.**
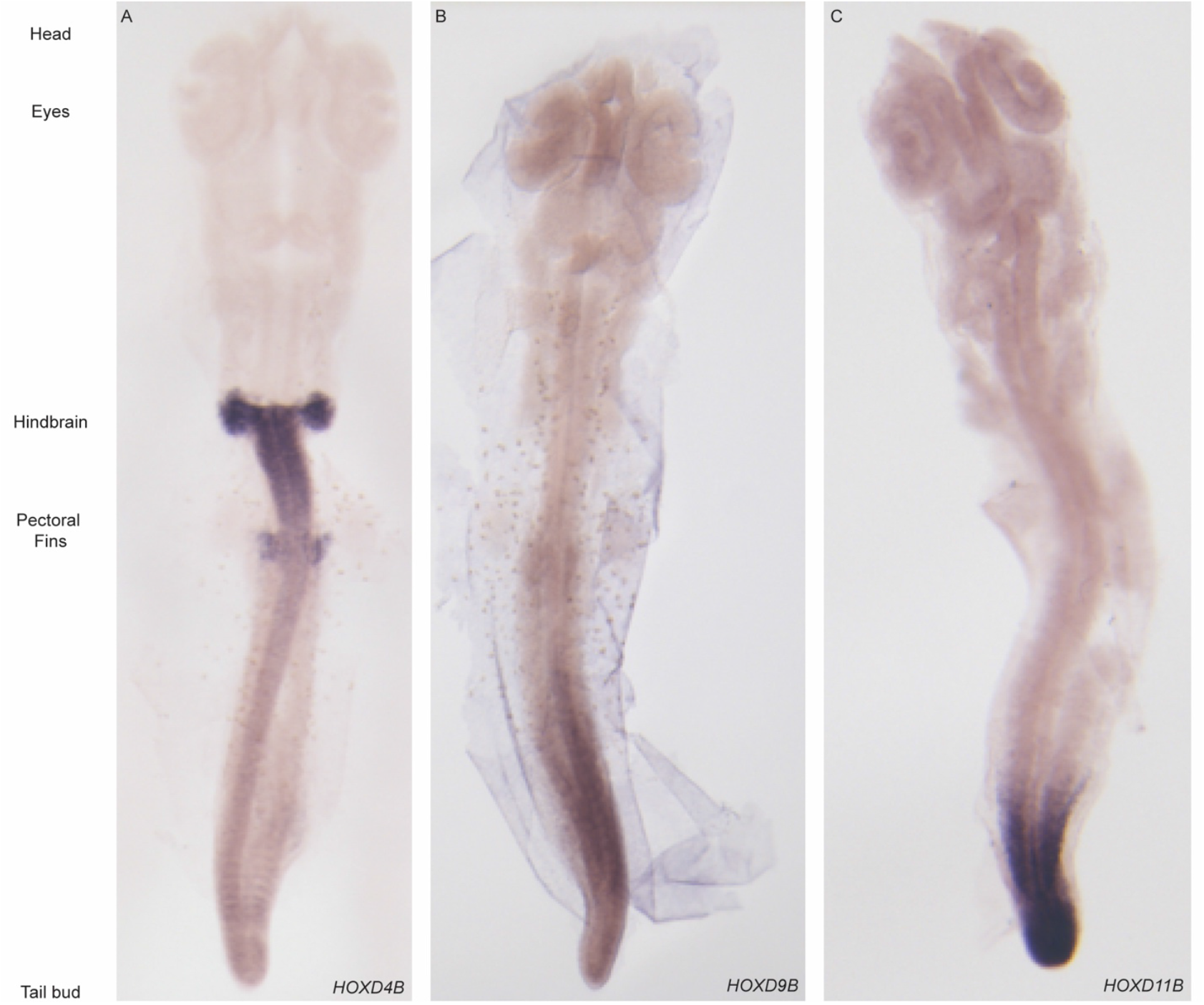
Embryonic expression of *Gasterosteus HOXDB* genes. *In situ* hybridization of *Gasterosteus aculeatus* embryos at Swarup stage 19/20 **A**. *HOXD4B*; **B**. *HOXD9B*; **C**. *HOXD11B*.

**Figure S3.**
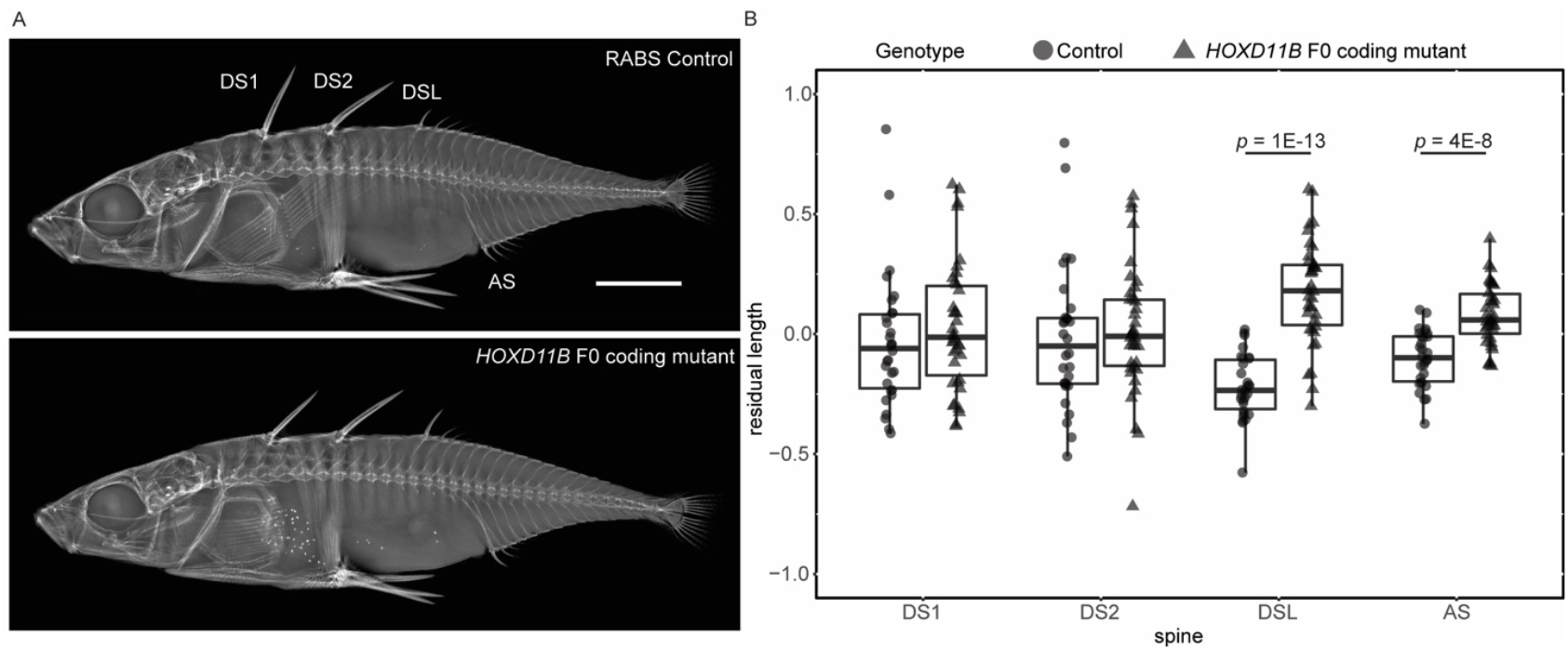
Coding mutations in *HOXD11B* cause length changes in *Gasterosteus* stickleback spines. **A**. X-rays of an uninjected sibling control Rabbit Slough (RABS) *Gasterosteus* (top) and a RABS *Gasterosteus* that was injected at the single cell stage with Cas9 and an sgRNA targeting the coding region of *HOXD11B* (bottom). Scale bar is 5mm. **B**. Quantification of spine length changes. DS1 and DS2 were not significantly different between controls and *HOXD11B* mutant fish. DSL and AS were significantly longer in the F0 mutants compared to the controls (two-tailed t-test; DSL p=1E-13, AS p=4E-08, n= 38 injected and n=30 control from 3 clutches combined). The y-axis is the residual after accounting for the standard length of fish.

**Figure S4.**
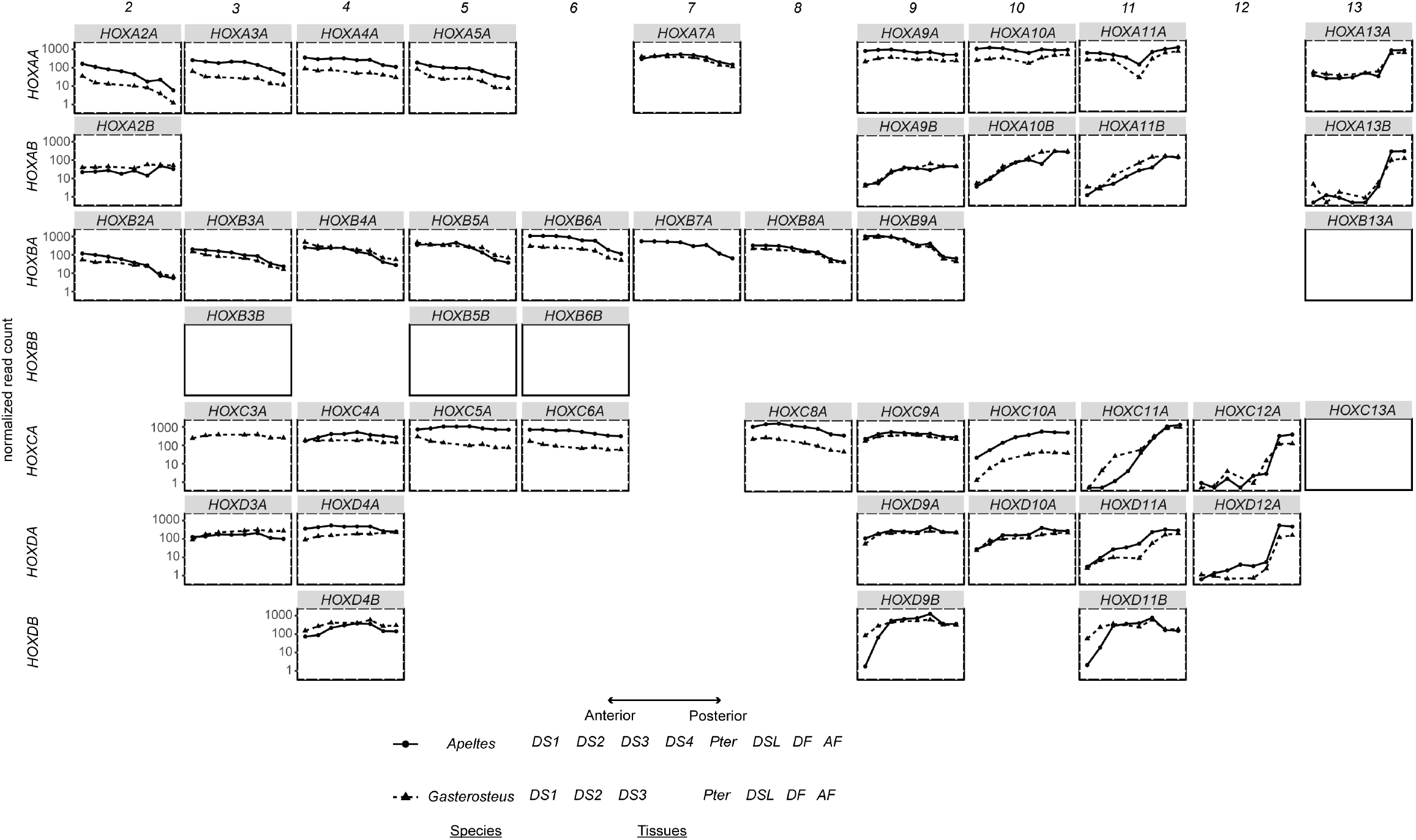
*Hox* gene expression patterns in *Gasterosteus* and *Apeltes* spines and fins. The expression patterns for each *Hox* gene in different stickleback *Hox* clusters are shown with normalized read count on the y-axis and tissue site on the x-axis. The tissues are organized by position from anterior to posterior along the dorsal side of the fish, with the anal fin at the end. The read count shown is the average across all samples for that species; the reads are normalized within each species but not between species. The genes with empty plots exist in both species but are not expressed in the tissues shown with the exception of *HOXB6B* and *HOXB7A*, which are located in a gap in the *Gasterosteus* assembly and thus was not scored; *HOXB6B* is not expressed in *Apeltes* but *HOXB7A* is expressed. *HOXA1A, HOXB1B*, and *HOXB1B* are present in the genomes but were not expressed and are not shown. The genes differentially expressed (padj < 0.01) between the largest anterior spines (DS1 and DS2) in *Gasterosteus* low-spine (three-spine) fish are *HOXA2A, HOXA5A, HOXA10B, HOXC3A, HOXC5A, HOXC9A, HOXC10A, HOXC11A, HOXD3A, HOXD4A, HOXD9A, HOXD10A, HOXD4B, HOXD9B*, and *HOXD11B*; the genes differentially expressed (padj < 0.01) between DS1 and DS3 in *Apeltes* low-spine (four-spine) fish are *HOXA10B, HOXC4A, HOXC8A, HOXC9A, HOXC10A, HOXD9A, HOXD10A, HOXD11A, HOXD4B, HOXD9B*, and *HOXD11B*.

**Supplemental Figure S5.**
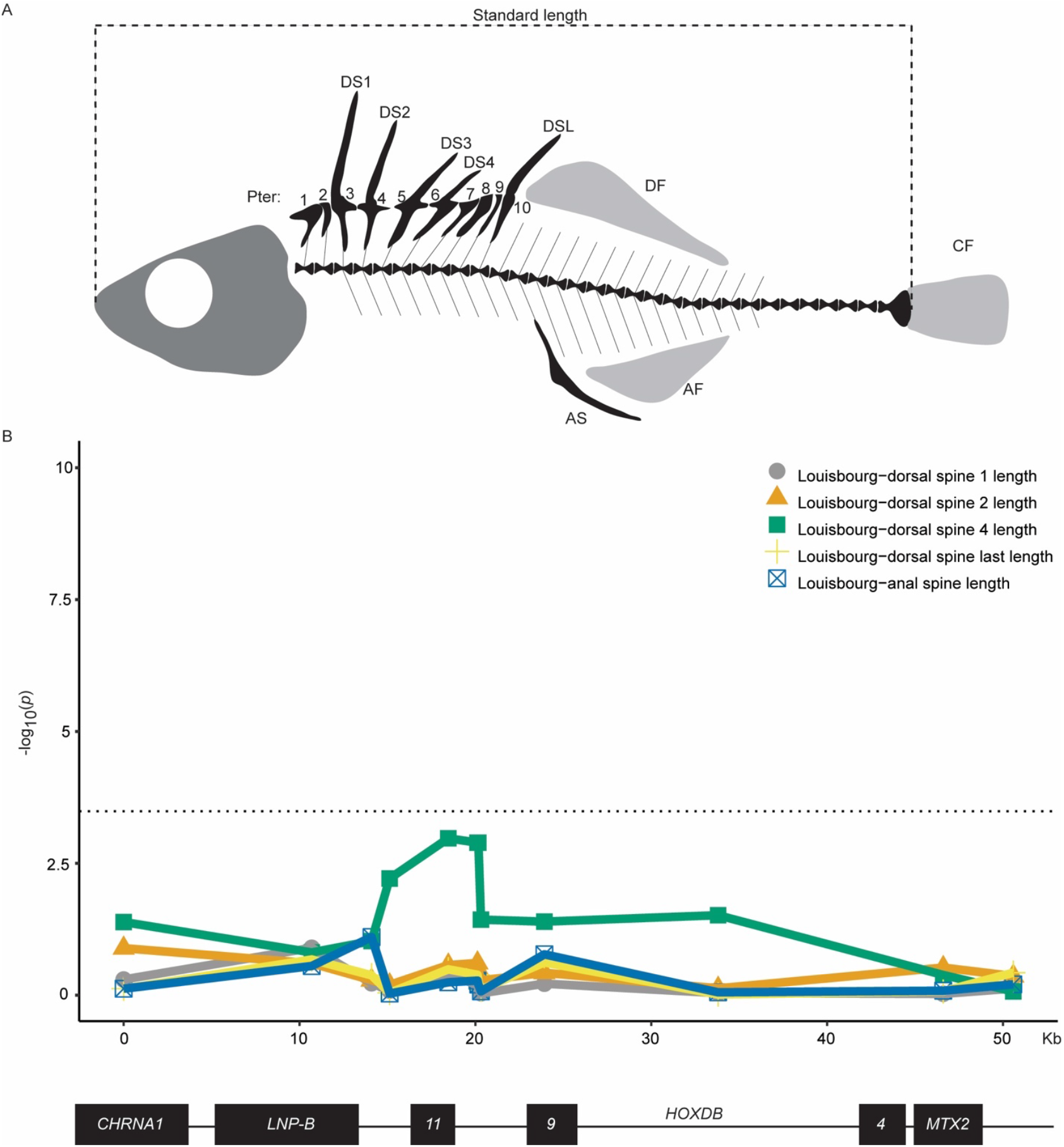
Spine anatomy and trait association mapping in Louisbourg *Apeltes*. **A**. Schematic of anatomical structures in an *Apeltes* fish with five dorsal spines. Typical A-P midline pattern: two non-spine bearing pterygiophores (Pter 1 and 2), dorsal spine 1 (DS1) on pterygiophore 3 (Pter3), dorsal spine 2 (DS2) on pterygiophore 4 (Pter4), dorsal spine 3 (DS3) on pterygiophore 5 (Pter5), dorsal spine 4 (DS4) on pterygiophore 6 (Pter6), three non-spine bearing pterygiophores (Pter7-9), and dorsal spine last (DSL) on pterygiophore 10 (Pter10). The three unpaired fins are shown in light gray: dorsal fin (DF), caudal fin (CF), and anal fin (AF). The anal spine (AS) is indicated on the ventral side of the fish. The standard length shown with the dotted line is from the anterior tip of the jaw to the posterior of the hypural plates. **B**. The association between *HOXDB* genotypes and length of DS1, DS2, DS4, DSL, and AS were not statistically significant. For significant results with DS3 length, see Figure 5.

**Figure S6.**
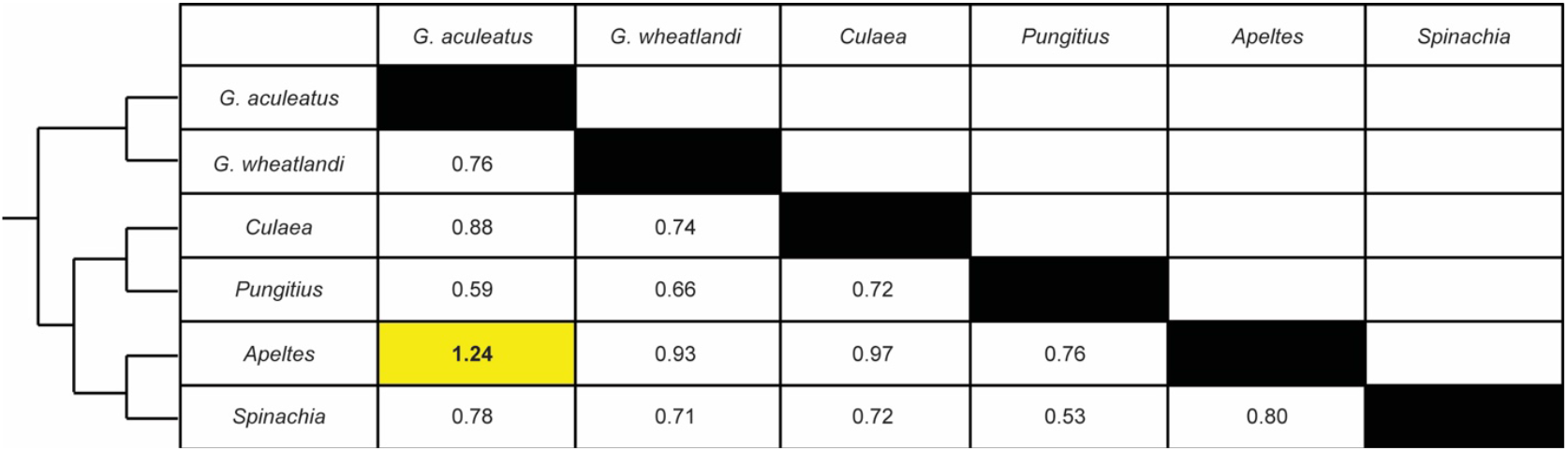
dN/dS values for *HOXD11B* between pairs of stickleback species. The tree on the left shows phylogenetic relationships of extant stickleback species (branch length not drawn to scale, (Kawahara et al., 2009; Liu et al., 2021)). The rate of non-synonymous to synonymous substitutions in *HOXD11B* is higher than 1 for *Gasterosteus* and *Apeltes* comparisons (yellow shading).

**Supplemental Figure S7.**
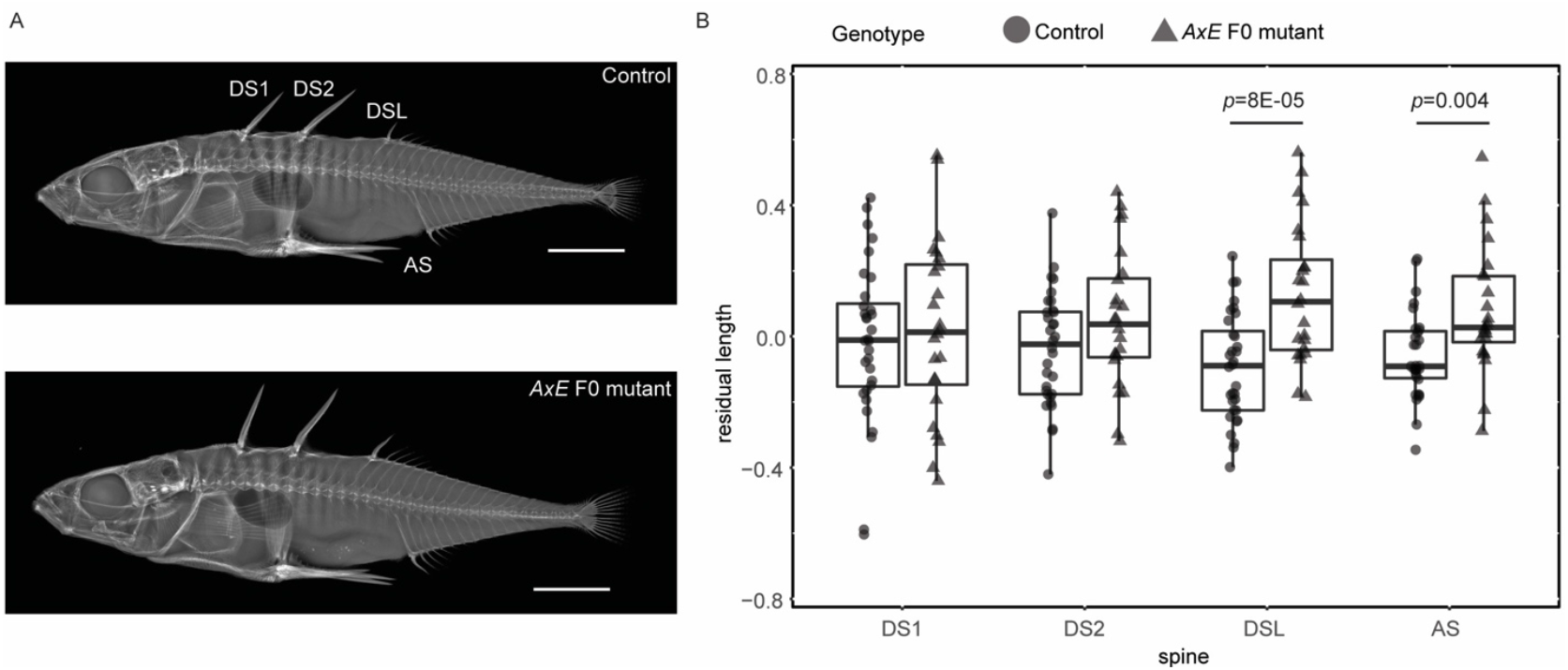
Mutation of *AxE* in a second anadromous *Gasterosteus* population also causes length changes in DSL and AS. **A**. Representative uninjected Little Campbell River sibling control fish (top) and injected *AxE* F0 mutant (bottom). Scale bar is 5 mm. **B**. Quantification of spine length difference. The residual after adjusting for standard length is on the y-axis and the spines ordered from anterior to posterior are on the x-axis. DS1 and DS2 do not show a significant difference in length between control and injected. DSL and AS were significantly longer in the injected compared to the control.

## Materials and Methods

### Stickleback care

Wild sticklebacks were captured using minnow traps, dip nets, or small minnow seines. The populations used for this study and their GPS coordinates are listed in Table S1. All sticklebacks were treated in accordance with the recommendations in the Guide for the Care and Use of Laboratory Animals of the National Institutes of Health, using protocols approved by the Institutional Animal Care and Use Committee of Stanford University (IACUC protocol #13834), in animal facilities accredited by the Association for Assessment and Accreditation of Laboratory Animal Care International (AAALAC).

### DNA extractions

DNA was isolated from single fins by placing them in lysis buffer (10mM Tris pH 8, 100mM NaCl, 10 mM EDTA, 0.5% SDS) with Proteinase K (333µg/mL, NEB P8107S) at 55°C for between four hours and overnight. DNA was extracted with Phenol:Cholorform:Isoamyl Alcohol 25:24:1 (Sigma, P3803) in phase lock tubes (Qiagen MaXtract High Density, 129056) and ethanol precipitated overnight. DNA was resuspended in TE Buffer (10mM Tris, 1 mM EDTA pH 8).

### QTL Mapping

A wild-caught female from Boulton Lake, British Columbia, Canada (BOUL) was crossed by *in vitro* fertilization to a marine male stickleback from Bodega Bay, California, USA (BDGB). F1 progeny were raised to adulthood in the laboratory in 30-gallon aquariums in RO-purified water with 3.5 ppt Instant Ocean salt and intercrossed to generate multiple F2 families. Sperm from F1 males was cryopreserved (Aoki et al., 1997), so single males could be crossed multiple times. F2 progeny fish were raised in lab for one year, then euthanized with 200 mg/L tricaine (Tricaine methanesulfonate, ANAD#200-226, Western Chemical Inc.) buffered to pH 7 with sodium bicarbonate and preserved in 70% ethanol.

DNA from each fish was extracted as described above, and the fish were genotyped using an Illumina GoldenGate genotyping array with 1536 features (Jones *et al*., 2012). Intensity data were processed using GenomeStudio 2011. Genotype clusters were inspected and adjusted manually, and uninformative or low-intensity SNPs were excluded from downstream analysis. Phasing and linkage map construction were performed with TMAP (Cartwright et al., 2007). The linkage map and phased genotype data were then loaded into R/qtl (Broman et al., 2003) and filtered to remove fish with fewer than 600 genotype calls and markers with fewer than 300 calls. A final map was generated with 343 F2s and 452 markers.

*Gasterosteus* anatomical traits and landmarks are diagrammed in Figure S1. Abdominal, caudal, and total vertebral counts, as well as pterygiophore number were counted from X-rays taken on a Faxitron UltraFocus X-ray cabinet (settings: 38 kV, 4.8 seconds). The lengths of dorsal spine 1, 2, and last and anal spine were measured based on the x-rays using Fiji (Schindelin et al., 2012). The measurements were adjusted by taking the residuals from multiple regression against standard length and sex. The presence or absence of a fourth spine and number of pterygiophores (six or more than six) were coded as binary traits (0 or 1). These phenotypes were used for QTL analysis in R/qtl using Haley–Knott regression via the “scanone” function, with a normal model for the length traits and a binary model for the spine and pterygiophore number (Broman et al. 2003). For the vertebral counts, a non-parametric (NP) scanone analysis was done. Permutation tests (*n* = 1,000) were used to establish LOD significance thresholds (*α* = 0.05) for each trait. The analysis is based on 340 F2 fish and a set of 452 SNP markers.

### *In situ* hybridization probes

RNA was extracted by homogenizing ten to twenty stage 19/20 embryos in Trizol using a FastPrep-24 machine (MP Biomedicals) and lysing matrix M. RNA was washed once with chloroform, precipitated with isopropanol, and resuspended in DEPC water. RNA was treated with on-column DNase and was cleaned up using QIAGEN RNeasy Mini (Qiagen, 74104) cleanup protocol. cDNA was made with the SuperScript(tm) VILO(tm) cDNA Synthesis Kit (Thermo Fisher, 11754050). For each riboprobe, RT-PCR amplification was done using the *in-situ* probe primers shown in Table S2. The *HOXD11B* probe was cloned into pCR2.1 TOPO (Invitrogen, K450001) in both orientations, and the *HOXD4B* and *HOXD9B* probes were cloned into pCRII-Blunt II-TOPO (Invitrogen, K280020) in both orientations. The vectors containing probe sequences were linearized with *BamHI* (Thermo Scientific, FD0054), and the sense and antisense probes were *in vitro* transcribed with T7 RNA Polymerase (Promega, P2075).

### Whole mount *in situ* hybridization

To determine the expression patterns of the *HOXDB* genes, whole mount *in situ* hybridizations at Swarup stages 19-20 were done as described by (Thisse and Thisse, 2008) with the following modifications. Embryos were manually dechorionated with two Dumont #5 - Fine Forceps (FST, 11251-10) after overnight fixation in 4% paraformaldehyde in PBS. To remove the pigmentation, they were bleached for ten minutes in 0.8% KOH, 3% hydrogen peroxide (30%), and 0.1% Tween20. Finally, embryos were permeabilized with Proteinase-K (NEB, P8107S) for ten seconds at 10 µg/ml in PBS with 0.1% Tween20.

### GFP knock-in

CRISPR-Cas9 was used to generate GFP reporter lines for *HOXD11B*, as described (Kimura et al., 2014). Cas9 protein (QB3 MacroLab University of California-Berkeley), a donor plasmid (pTia1l-hspGFP, deposited at Addgene, containing hsp70, GFP, and a sgRNA target site), and two sgRNAs were injected. One sgRNA (*HOXD11B*-GFP-sgRNA, Table S3) targeted the region 346bp upstream of the endogenous *HOXD11B* start codon, and one targeted the donor plasmid. The *HOXD11B*-GFP-sgRNA was designed as previously described (Wucherpfennig et al., 2019). Tia1l sgRNA was used to cut the plasmid (Lackner et al., 2015) and has a sequence not present in the *Gasterosteus aculeatus* genome. The injection mix contained a final concentration of 1 μg/μl Cas9 protein, 31 ng/μl of Tia1l sgRNA, 31 ng/μl of the *HOXD11B*-GFP-sgRNA, 0.05% phenol red, and was adjusted to the final concentration with 10mM Tris pH 7.5.

Fertilized eggs from *Gasterosteus* Little Campbell River (LITC) fish were injected at the single cell stage, and embryos were screened at st20 (~84 hpf) for GFP expression. Embryos with GFP expression were raised to stages 29-31 (18dpf) and imaged again. The fry were anesthetized with 3 mg/L tricaine (Tricaine methanesulfonate, ANAD#200-226, Western Chemical Inc.). Imaging was done with a MZFLIII fluorescent microscope (Leica Microsystems, Bannockburn, IL) using GFP2 filters and a ProgResCF camera (Jenoptik AG, Jena, Germany). GFP positive fish were grown to adulthood and crossed to wild-type LITC fish once they reach sexual maturity at approximately seven months of age. Progeny embryos were screened at st20 (~84 hpf) for GFP expression, and GFP-positive fish were raised to adulthood. To confirm integration and orientation of the GFP construct, primers were designed upstream and downstream of the *HOXD11B*-GFP-sgRNA site and in the plasmid on either side of the sgRNA cut site within the hsp70 promoter or the TOPO backbone (Table S2). All combinations of primers were tested by PCR; the presence and absence of bands was used to determine the orientation. Sanger sequencing of those products was used to determine the exact site of integration and any resulting deletions.

### Generation of *HOXD11B* coding and regulatory mutations using CRISPR-Cas9

Mutations in the coding regions of *HOXD11B*, were generated by injecting Cas9 protein and an sgRNA targeting the first exon after the start codon (*HOXD11B-*coding-sgRNA, Table S3). The sgRNA was designed and synthesized as previously described (Wucherpfennig et al., 2019). The injection mix included 1 μg/μl of Cas9 protein, 300 ng/μl of the sgRNA, 0.05% phenol red, and was adjusted to final concentration with 10mM Tris pH 7.5. This mix was injected into fertilized eggs from two anadromous *Gasterosteus* populations (Little Campbell River, British Columbia, Canada and Rabbit Slough, Alaska, USA) at the single cell stage. Mutations were confirmed by PCR (using Phusion High-Fidelity DNA Polymerase (Thermo Scientific, F-530L), GC buffer, and 3% DMSO) with *HOXD11B*-coding_1F and 1R and *HOXD11B*-coding_1F and 1R (Table S2). The PCR program was 98°C (3 min), then 35 cycles at 98°C (10 s) / 60°C (30 s) / 72°C (30 s), and a final extension at 72°C for 10 min.

Two different strategies were used to delete *AxE*, the conserved enhancer (466 bp) between *HOXD9B* and *HOXD11B*: 1) three sgRNAs (*AxE*-sgRNA_1, 2, and 3, Table S3) and a 60bp repair phosphorothioate modified oligo (IDT) with 30bp of homology to either side of the enhancer (Renaud et al., 2016); or 2) a total of six sgRNAs (*AxE*-sgRNA_1 through 6, Table S3) targeting the edges and middle of the enhancer. The sequence for the repair oligo was G*A*A CGT AAA AGG ATT CAG GAG CTC AAG CGA GTC GGT TCC AAA CGT GTC GTT GCC CAG C*A*G with the asterisks indicating the phosphorothioate bonds. If the first two bases of the sgRNA target sequences were not Gs, then they were replaced to aid in the transcription of the sgRNA. The injection mix included 1 μg/μl of Cas9 protein, 300 ng/μl of total of the sgRNAs (100 ng/μl of each for strategy 1, 50 ng/μl of each for strategy 2), 1.5 pmol/μl repair oligo (strategy 2 only), 300mM KCl (Burger et al., 2016), 0.05% phenol red, and was adjusted to final concentration with water. The mutations were confirmed by PCR as described above, except that the extension time was increased to one minute and the annealing temperature was 64°C. Two sets of primers were used, with the first amplifying only the 571 bp region including the enhancer and the second including ~3.6 kb around the enhancer to identify potential larger mutations (Table S2).

### *Apeltes quadracus* association mapping

Fourspine sticklebacks (*Apeltes quadracus*) were collected in May 2018 and May and July 2019 using minnow traps and dip nets from Fortress Louisbourg (Site 325) and Tidnish River Site 3 (Site 171) (Blouw, 1982) (GPS coordinates in Table S1). Sticklebacks were euthanized as described above and were fixed in 70% ethanol or Alfred Lamb’s Navy Dark Rum 151 Proof. *Apeltes* anatomical traits and landmarks are diagrammed in Figure S5. Fish were phenotyped for spine number using a Faxitron UltraFocus X-ray cabinet. The dorsal and anal spine lengths and standard length of the Louisbourg fish were measured in triplicate using digital calipers, and the average of the measurements was used as the length. The residuals were calculated for each spine length taking into account the standard length of the fish. Pectoral and caudal fins were clipped to make genomic DNA as described above.

To identify potential genotyping markers (microsatellites, indels, or SNPs) without a reference genome for *Apeltes, HOXDB Apeltes* sequence was amplified by PCR using primers (*PUNG-GAC*_1-11, Table S2) conserved between the *Gasterosteus aculeatus* (Jones et al., 2012b) and *Pungitius* genomes (GenBank assembly accession numbers: GCA_003399555.1, GCA_003935095.1, GCA_902500615.2) (Nelson and Cresko, 2018; Varadharajan et al., 2019) (Table S2). The PCR products were then TOPO cloned into pCRII-Blunt II-TOPO (Invitrogen, K280020), miniprepped, and Sanger sequenced from two to four individuals with differing spine numbers to identify variable regions.

The regions between the PCR products around *LUNAPARK-B, HOXD11B*, and *HOXD9B* were filled by designing primers spanning the existing products (Table S2). These PCR products were also cloned as described above and Sanger sequenced. Additional internal sequencing primers were designed to fully sequence the products (Table S2).

Twelve markers were identified throughout the *Apeltes HOXDB* cluster and were scored in 211 fish from Louisbourg Fortress (7 six-spine, 99 five-spine, 104 four-spine, 1 three-spine) and 121 fish from Tidnish River 3 (1 six-spine, 59 five-spine, 59 four-spine, 1 three-spine, 1 two-spine).

Microsatellite markers were amplified using the universal fluorescent primer system described by (Schuelke, 2000). A 20 µl PCR reaction mixture contained 2x Master Mix (Thermo Fisher, K0171), 0.5 µM 6FAM M13 Forward universal primer, 0.125 µM forward primer, 0.5 µM reverse primer, and 10 ng of genomic DNA. The PCR program was 94°C (5 min), then 30 cycles at 94°C (30 s) / 58°C (45 s) / 72°C (45 s), followed by 8 cycles 94°C (30 s) / 53°C (45 s) / 72°C (45 s), and a final extension at 72°C for 10 min. For *AQ-HOXDB*_2, the cycle number was reduced from 30 to 27. The PCR was cleaned up using ExoSAP-IT PCR Product Cleanup Reagent (Applied Biosystems, 78205.1.ML), and the fragment sizes were analyzed on Applied Biosystems 3730xl Genetic Analyzer. Peaks were called using the Microsatellite plugin for Geneious.

The PCR reaction mix for indel and SNP markers was 2x PCR Master Mix, 0.5 µM Forward Primer, 0.5 µM Reverse Primer and 10 ng of genomic DNA. The PCR program was 95°C (5 min), then 35 cycles at 95°C (30 s) / 54°C (45 s) / 72°C (30 s), and a final extension at 72°C for 10 min. The one exception was *AQ-HOXDB*_6, where the PCR was done with Phusion High-Fidelity DNA Polymerase (Thermo Scientific, F-530L), GC buffer, and 3% DMSO, and the PCR program was 98°C (3 min), then 35 cycles at 98°C (10 s) / 60°C (30 s) / 72°C (10 s), and a final extension at 72°C for 10 min. The *AQ-HOXDB*_5 PCR product was digested with *BssSI-*v2 (NEB, R0680L), and the *AQ-HOXDB*_6 PCR product was digested with *NdeI* (Thermo Scientific FD0583). PCR products were run on a 2% agarose gel to detect size differences.

The allele frequencies in low-spine (two-to four-spine) and high-spine (five-to six-spine) fish were compared using CLUMP (Sham and Curtis, 1995), which performs a modified chi-square analysis to determine significance of allele frequency differences. For microsatellite markers, the negative log p-values of the chi-squared value (T4) from the 2×2 contingency table generated by CLUMP are shown in Figure 5. For indel or SNP markers, a chi-square test was performed in R (v. 3.6.1). The association between spine length (the average of triplicate measurements) and genotype was quantified using an ANOVA performed in R, using the residual spine length after accounting for the standard length of the fish.

### *Apeltes* genome assembly

Whole genome sequencing using 10X Genomics chromium linked read technology was performed on two *Apeltes quadracus* from the Louisbourg Fortress population (one four-spine and one five-spine). Genomic DNA was extracted from the brains of the fish and prepared using Qiagen MagAttract HMW DNA kit. The linked-read data of each fish were assembled using Supernova v.2.1.1 with default settings (Weisenfeld et al., 2014). The 4-spine *Apeltes* assembly had 16,216 scaffolds with 416,290,932 bases (scaffold N50: 393,888 bp; L50: 247; N90: 7,174 bp; L90: 3,684). The 5-spine *Apeltes* assembly had 24,175 scaffolds with 397,678,333 bases (scaffold N50: 69,128 bp; L50:1,629 ; N90: 4,805 bp; L90: 10,192).

To be able to use GATK in the allele-specific RNA-sequencing pipeline, the genome needed to be on fewer scaffolds than were generated by the linked read data. To achieve this, we started with the 4-spine *Apeltes* assembly and assumed that the chromosome structure of *Apeltes* and *Gasterosteus* are similar. We used a reference guided scaffold approach by generating global genome to genome alignments with minimap2 (Li, 2018) and MUMmer (Marçais et al., 2018). The alignment information was processed by RaGOO (Alonge et al., 2019) to order and orient contigs into scaffolds, which resulted in the *Apeltes* genome reference used in the GATK allele-specific RNA-sequencing pipeline.

### High-spine *Gasterosteus* genome assembly

Whole genome sequencing using 10X Genomics chromium linked read technology was performed on two four-spine *Gasterosteus aculeatus* from the F5 generation of the BOUL-BDGB QTL cross. Genomic DNA was extracted from the brains of the fish and prepared using Qiagen MagAttract HMW DNA kit. The linked-read data of each fish were assembled using Supernova v.2.1.1 with default settings (Weisenfeld et al., 2014).

Whole genome sequencing using PacBio HiFi technology was also performed on one four-spine *Gasterosteus aculeatus* from the F5 generation of the BOUL-BDGB QTL cross. Genomic DNA was extracted from the testes of the fish and prepared using Qiagen MagAttract HMW DNA kit. The genome was assembled using CANU. The purge haplotigs pipeline (https://bitbucket.org/mroachawri/purge_haplotigs/src/master/) was used to phase the alleles and identify the contigs that appeared twice in the assembly. The final assembly had 483 scaffolds with a total of 489,328,730 bases (scaffold N50: 3,689,351 bp; L50: 37; N90: 633,554 bp; L90:166).

### Transgenic enhancer assays

To identify and confirm sequence variants in the intergenic region between *HOXD9B* and *HOXD11B*, the ~6 kb intergenic region from *Apeltes* was amplified from a three-spine and a six-spine *Apeltes* from the Louisbourg Fortress population and a three-spine and a six-spine from the Tidnish River 3 population with Phusion High-Fidelity DNA Polymerase (Thermo Scientific, F-530L) in GC Buffer and 3% DMSO using primers in Table S2. The resulting products were TOPO cloned into pCRII-Blunt II-TOPO. Colonies were miniprepped and Sanger sequenced. To generate the plasmids for the enhancer assay, the low- and high-spine versions of the ~600 bp region that contains *AxE* were then amplified with primers that included overhangs homologous to the PT2HE GFP reporter vector (Howes et al., 2017; Kotani et al., 2006). The reporter vector was cut with *EcoRV* (Thermo Scientific, ER0201) and the insert and vector were joined using Gibson Cloning (NEB # E2611S). The resulting plasmids were screened by *SacI* (Thermo Scientific, ER1131) restriction digest and further Sanger sequenced to check for mutations. The 587 bp high-spine and 611 bp low-spine *Apeltes AxE* sequences are available in GenBank at OKxxxxxx and OKxxxxxx, respectively.

Transgenic *G. aculeatus* sticklebacks were generated by microinjection of fertilized eggs at the single cell stage from LITC *G. aculeatus* as described in (Chan et al., 2010). Plasmids (25 ng/µl) were injected with Tol2 transposase mRNA (36 ng/µl) and 0.1% Phenol red as described in (Hosemann et al., 2004). Tol2 mRNA was synthesized by *in vitro* transcription using the mMessage mMachine SP6 kit (Invitrogen, AM1340) from pCS-TP plasmid (Kawakami et al., 2004) cut with *Bsp120I* (Thermo Scientific, ER0131). Transgenics were imaged at st20 (~84 hpf) and st29/31 (~18-30 dpf) as described in the GFP knock-in section above. The hsp70 promoter drives expression in the lens of the eye by nine days post fertilization (Nagayoshi et al., 2008). At st29/31, bilateral lens GFP expression was used to identify less mosaic fish.

### dN-dS calculation

dN-dS calculations were performed in R using ape v5.3 (Paradis and Schliep, 2019). The sequence alignments for each gene (*HOXD11B, HOXD9B*, and *HOXD4B*) were generated in Geneious using translation alignment. The mature transcript for each gene was determined based on the splicing that has been validated using cDNA in *G. aculeatus*. The sequences for *P. pungitius* were determined by BLASTN (Altschul et al., 1990) of the *G. aculeatus* exons against the genome (*Pungitius*: GCA_003935095.1 (Nelson and Cresko, 2018)). The sequences for *Apeltes* were identified from our genome assembly. The sequences for *Gasterosteus wheatlandi, Culaea inconstans*, and *Spinachia spinachia* were identified by BLAST of the *G. aculeatus* exons against unassembled short reads from whole genome sequencing of the respective species ((Liu et al., 2021) and Catherine Peichel, personal communication).

### RNA-sequencing

For *Gasterosteus* RNA-sequencing, a lab-raised Little Campbell River anadromous female with three dorsal spines was crossed to a high-spine *Gasterosteus* male with five dorsal spines. The high-spine *Gasterosteus* line is the F5 generation of the original QTL cross between BOUL and BDGB used to identify the *HOXDB* locus. The fish have been selected for high spine number, and by F5, more than 80% of fish have four or more dorsal spines. To confirm that the fish carried the BOUL allele at the *HOXDB* locus, the allele was amplified using BOUL-*HOXDB*_1F and 1R (Table S2) with Phusion High-Fidelity DNA Polymerase (Thermo Scientific, F-530L) in GC Buffer and 3% DMSO using a 2-step PCR program (94°C (1 min), then 30 cycles at 98°C (10 s) / 68°C (15 minutes), and a final extension at 72°C for 10 min) and run on an agarose gel. The Boulton allele is ~15kb, and the Bodega Bay allele is ~1.9 kb. The sequences of the two alleles are available at OKxxxxxx (Bodega Bay) and OKxxxxxx (Boulton) in GenBank.

The resulting clutch was raised to 11-13mm. The fry were euthanized in 200 mg/L tricaine buffered to pH 7 with sodium bicarbonate. The fish were dissected on a 2% agarose plate with size 00 insect pins and Vannas Spring Scissors - 2.5mm Cutting Edge (FST, 15000-08). The tissues (shown in Figure 4A were: dorsal spine 1, dorsal spine 2, dorsal spine 3, dorsal spine last, blank pterygiophore, dorsal fin, and anal fin) were flash frozen in liquid nitrogen in FastPrep Tubes (MP Biomedicals, MP115076200). DNA was extracted from tails and genotyped to ensure fish had informative SNPs in the coding region of *HOXD11B* and *HOXD9B*. For *HOXD11B*, the primers were *HOXD11B*-coding_1F and 1R (Table S2). For *HOXD9B*, the primers were *HOXD9B*-coding_1F and 1R (Table S2). The PCR conditions were the same as described above for the confirmation of *HOXD11B* CRISPR mutants.

Based on the genotyping, twelve three-spine progeny and six four-spine progeny were chosen for RNA extraction, library prep, and sequencing. Samples for RNA extraction were homogenized using MP FastPrep 2 × 20 seconds with Matrix M with a five-minute rest in between. RNA extractions were performed using NucleoSpin® RNA XS (Takara) and resuspended in 20ul of RNase free water. RNA was quantified by Qubit (Invitrogen) using HS Assay Kit (Invitrogen, Q32851). A subset of samples was quality controlled to check the RIN values by Bioanalyzer using the RNA 6000 Pico Kit (Agilent, 5067-1513). The RINs were between 8.2 and 10, with most higher than 9.6. Sequencing libraries were generated with Illumina Stranded mRNA Prep kit (Illumina, 20040532) and 20-100 ng of RNA (depending on the amount of RNA; if less than 100 ng was extracted, the entire sample was used). The PCR cycle number was determined by qPCR and was generally: 12 cycles for embryo samples with 200 ng of RNA, 14 cycles for dorsal spine and pterygiophore samples with 100 ng of RNA, 13 cycles for dorsal and anal fin samples with 100 ng of RNA, and 15 cycles for samples with less than 100 ng of RNA input. Quality control of libraries was done by Qubit with a dsDNA HS Assay Kit (Invitrogen, Q32851) to check the concentration and by BioAnalyzer with a high sensitivity kit (Agilent, 5067-4626) to check the size. Libraries were sequenced to a coverage of ~30 million reads on a NovaSeq 6000 (2 × 150 bp) by NovoGene. Reads were trimmed with Cutadapt (Martin, 2011) using the TrimGalore wrapper (https://github.com/FelixKrueger/TrimGalore), and reads were mapped to the *gasAcu1-4* reference genome (https://datadryad.org/stash/dataset/doi:10.5061/dryad.547d7wm6t) with STAR two-pass mapping (Dobin et al., 2013). For allele-specific expression analysis, the base quality was adjusted and variants were called using GATK as recommended by Broad Institute best practices (Van der Auwera et al., 2013; Depristo et al., 2011). The reads at each site were counted using GATK ASEReadCounter. We required that SNPs be called as heterozygous in at least one tissue of each fish, that the number of reads at a given site be greater than 12 (three-spine) or 14 (four-spine) for each fish, and that the overall minor allele frequency be greater than 5%. To quantify the allele-specific expression differences seen between the dorsal tissues and the anal fin (control), we took the log2 ratio of the reference reads to the alternate reads within each sample and compared each dorsal tissue to the anal fin using a Mann-Whiney U test.

To improve the gene predictions and recover any novel transcripts for differential gene expression analysis, StringTie was used along with the existing Ensembl annotations to predict the transcripts (Pertea et al., 2015). Given the large number of reads, bam files were filtered by quality, downsampled to 20%, and merged into one file that was used as the input for StringTie. The merge function was used to add back in genes from the Ensembl annotations not present in the sequenced samples. All *Hox* genes were manually checked. In some cases, the two genes were merged into one due to their close proximity; these were manually separated in the GTF file. FeatureCounts was then used with the new GTF file to assign reads to genes (Liao et al., 2014). Differential gene expression between different tissues was performed in DESeq2 (Love et al., 2014).

For *Apeltes* RNA-sequencing, the same protocol was followed as detailed for *Gasterosteus* above, with the following differences. The spines, blank pterygiophore, dorsal fin, and anal fin were dissected from Louisbourg *Apeltes* clutches raised in the lab to 11-13mm. The fry were genotyped for the two peak association mapping marker (*AQ-HOXDB*_6 and 7) using the same primers and conditions described above under *Apeltes* association mapping. For allele-specific expression analysis, four fish with L/LHR (heterozygous for the 18bp indel allele (*AQ-HOXDB*_7) and homozygous for the 2 adjacent SNPs, GG (*AQ-HOXDB*_6)) genotype, four fish with H/L (heterozygous for the 18bp indel and the 2 adjacent SNPs) genotype, and three fish with H/LHR (homozygous for the 18bp deletion and heterozygous for the 2 adjacent SNPs) genotype were sequenced. Three four-spine L/L (homozygous for the 18bp intact allele and homozygous for the 2 adjacent SNPs, GG) genotype were also sequenced to examine the expression differences between tissues. To generate gene predictions for the *Apeltes* genome, StringTie was used; the *Hox* genes were identified by BLAST and manually named in the GTF file (Altschul et al., 1990). For the high-spine fish, allele-specific analysis was performed as described above for *Gasterosteus*. The Apeltes 10X linked read data was used as input for 10X Long Ranger (v. 2.2.2) to generate a vcf file of known variants for GATK Baserecalibator. Because the clutch size of *Apeltes* is smaller and thus the number of replicates was lower than in the *Gasterosteus* analysis, we use a Fisher’s Exact Test to compare expression in dorsal tissues to the anal fin. We summed the references and alternate reads within each tissue to generate a 2×2 contingency table. For the analysis show in Figure S4, differential gene expression between tissues was performed in DESeq2 (Love et al., 2014).

## Data and code availability

The raw and processed allele-specific RNA-sequencing data in this paper will be available in the NCBI GEO database: GSExxxxxx (subseries GSExxxxxx, GSExxxxxx, GSExxxxxx, GSExxxxxx). The PacBio HiFi and 10X linked read data from *Gasterosteus* high-spine sequencing will be available in the NCBI databases under BioProject number: PRJNAxxxxxx. The 10X linked read data from *Apeltes quadracus* four- and five-spine fish will be available under BioProject number: PRJNAxxxxxx. The sequence surrounding *AxE* in *Gasterosteus* from the two parental QTL populations and the *Apeltes AxE* sequences tested in transgenic assays will be available in GenBank (OKxxxxxx, OKxxxxxx, OKxxxxxx, OKxxxxxx). QTL mapping files, phenotype data files, association mapping genotype files, and code will be available at Mendeley Data. The pTia1l-hspGFP plasmid will be available from Addgene. Other materials will be made available upon request.

